# Crossover patterning through kinase-regulated condensation and coarsening of recombination nodules

**DOI:** 10.1101/2021.08.26.457865

**Authors:** Liangyu Zhang, Weston Stauffer, David Zwicker, Abby F. Dernburg

**Author notes:** Equal contribution (Liangyu Zhang, Weston Stauffer).

## Abstract

Meiotic recombination is highly regulated to ensure precise segregation of homologous chromosomes. Evidence from diverse organisms indicates that the synaptonemal complex (SC), which assembles between paired chromosomes, plays essential roles in crossover formation and patterning. Several additional “pro-crossover” proteins are also required for recombination intermediates to become crossovers. These typically form multiple foci or recombination nodules along SCs, and later accumulate at fewer, widely spaced sites. Here we report that in *C. elegans* CDK-2 is required to stabilize all crossover intermediates and stabilizes interactions among pro-crossover factors by phosphorylating MSH-5. Additionally, we show that the conserved RING domain proteins ZHP-3/4 diffuse along the SC and remain dynamic following their accumulation at recombination sites. Based on these and previous findings we propose a model in which recombination nodules arise through spatially restricted biomolecular condensation and then undergo a regulated coarsening process, resulting in crossover interference.

## Introduction

Crossover recombination during meiosis gives rise to genetic diversity and is essential for sexual reproduction. Genetic studies in Drosophila in the early 20th century first revealed the existence of “crossover interference,” the phenomenon that crossovers detected between the same pair of homologs are nonrandomly far apart (Muller, 1916; Sturtevant, 1915). Since these seminal findings, crossover interference has been detected in most organisms in which meiotic recombination has been studied, including plants, fungi, and metazoans. In many eukaryotes, including the nematode *C. elegans*, a single crossover per chromosome pair is the norm during each meiosis. Mutations that increase crossing-over often result in aneuploid gametes, indicating that the crossover number is tightly regulated to create an optimal number of physical links between homologs to ensure their segregation.

Crossovers arise through repair of meiotic double-strand breaks (DSBs). Only a subset of DSBs is repaired to form crossovers; most are repaired through alternate pathways that lead to noncrossover products (a.k.a. “simple gene conversions”). Additionally, only crossovers produced by a meiosis-specific “pro-crossover pathway,” a.k.a. “Class I” crossovers, are subject to interference. This widely conserved pro-crossover pathway is also referred to as the “ZMM” pathway, based on the names of several of the proteins involved in budding yeast (Lynn et al., 2007). Additional noninterfering “Class II” crossovers can arise in some organisms as byproducts of repair pathways that lead mostly to noncrossovers (reviewed by Gray and Cohen, 2016).

Double strand breaks (DSBs) are induced as chromosomes pair and synapse during early meiotic prophase. They are resected to produce 3’ single-stranded overhangs, which recruit single-strand binding proteins and strand-exchange factors, including Dmc1 and/or Rad51. These form protein-DNA filaments that can invade the homologous chromosome and form D-loops, then undergo further processing to form “joint molecules” (JMs), which have the potential to be resolved as either crossovers or noncrossovers. JMs recruit several pro-crossover factors, which collectively protect them from noncrossover resolution. Eventually these factors recruit crossover resolvases, leading to resolution and the formation of chiasmata, the cruciform structures that maintain connections between homologs until the first meiotic division.

In diverse organisms, pro-crossover factors assemble to form “early recombination nodules” (ERNs), ellipsoidal bodies that can be observed by transmission electron microscopy (Wettstein et al., 1984). ERNs are typically 25-100 nm in diameter, and do not show interference in their spatial distribution along chromosomes. As meiotic prophase progresses, ERNs are typically replaced by fewer, larger bodies termed “late recombination nodules” (LRNs). These do show interference and closely mirror the patterning of crossovers as measured through genetic assays. Thus, the patterning of late nodules underlies crossover interference, but the mechanisms controlling this patterning remain poorly understood and controversial.

ERNs and LRNs share most of their molecular components. In most organisms these include MutSɣ, a heterodimer of two meiosis-specific MutS homologs, Msh4/Msh5 heterodimer (HIM-14/MSH-5 in *C. elegans*) that bind directly to JMs (Snowden et al., 2004). Members of a widely conserved family of RING finger proteins are also associated with ERNs and LRNs and are required for their appearance. The MutLɣ complex (Mlh1/Mlh3) is specifically associated with LRNs in many plants, fungi, and mammals, and plays an essential role in resolution of intermediates to form crossovers (Anderson et al., 1999; Rogacheva et al., 2014). *C. elegans* lacks a Mlh3 homolog, as do many other nematodes, and *mlh-1* mutants are viable and fertile, indicating that the protein is dispensable for meiosis (Meier et al., 2018); it is not yet clear how resolvases are recruited to execute crossover resolution. Both early and late nodules are physically associated with the synaptonemal complex (SC), the protein matrix that assembles between homologous chromosomes during meiosis. 3-dimensional analysis by electron microscopy of serial sections has indicated that the nodules sit astride the SC rather than being embedded within it (Carpenter, 1975; Schmekel and Daneholt, 1998; Zickler, 1977; Zickler and Kleckner, 2015).

It is uncertain whether recombination nodules have been observed by EM in *C. elegans* (see Discussion). However, fluorescent labeling of pro-crossover proteins, particularly MSH-5, reveals multiple faint early foci along each SC, and eventually a single focus per SC that colocalizes with COSA-1, a cyclin homolog (Woglar and Villeneuve, 2018; Yokoo et al., 2012) Figure 1). These late foci correspond to sites of eventual crossovers, and their appearance is regarded as “crossover designation.” We consider the early and late fluorescent foci to be functionally equivalent to ERNs and LRNs. The molecular requirements for and composition of early and late foci are very similar, as detailed below. A heterodimeric complex comprised of two RING domain proteins, ZHP-3/ZHP-4, is required for all foci (Zhang et al., 2018). This complex initially localizes along the SC but eventually concentrates at the late foci. A related complex, ZHP-1/ZHP-2, is specifically required for the appearance of late foci, but these proteins remain distributed along the SC, rather than concentrating at foci. We recently reported that the transition from early to late foci in *C. elegans* is triggered by inactivation of CHK-2 kinase activity, which normally occurs at mid-pachytene (Zhang et al., 2021). Mutations that impair the formation of JMs along one or more chromosomes trigger a crossover assurance checkpoint, which delays the timing of CHK-2 inactivation and crossover designation (Yu et al., 2016).

**Figure 1.**
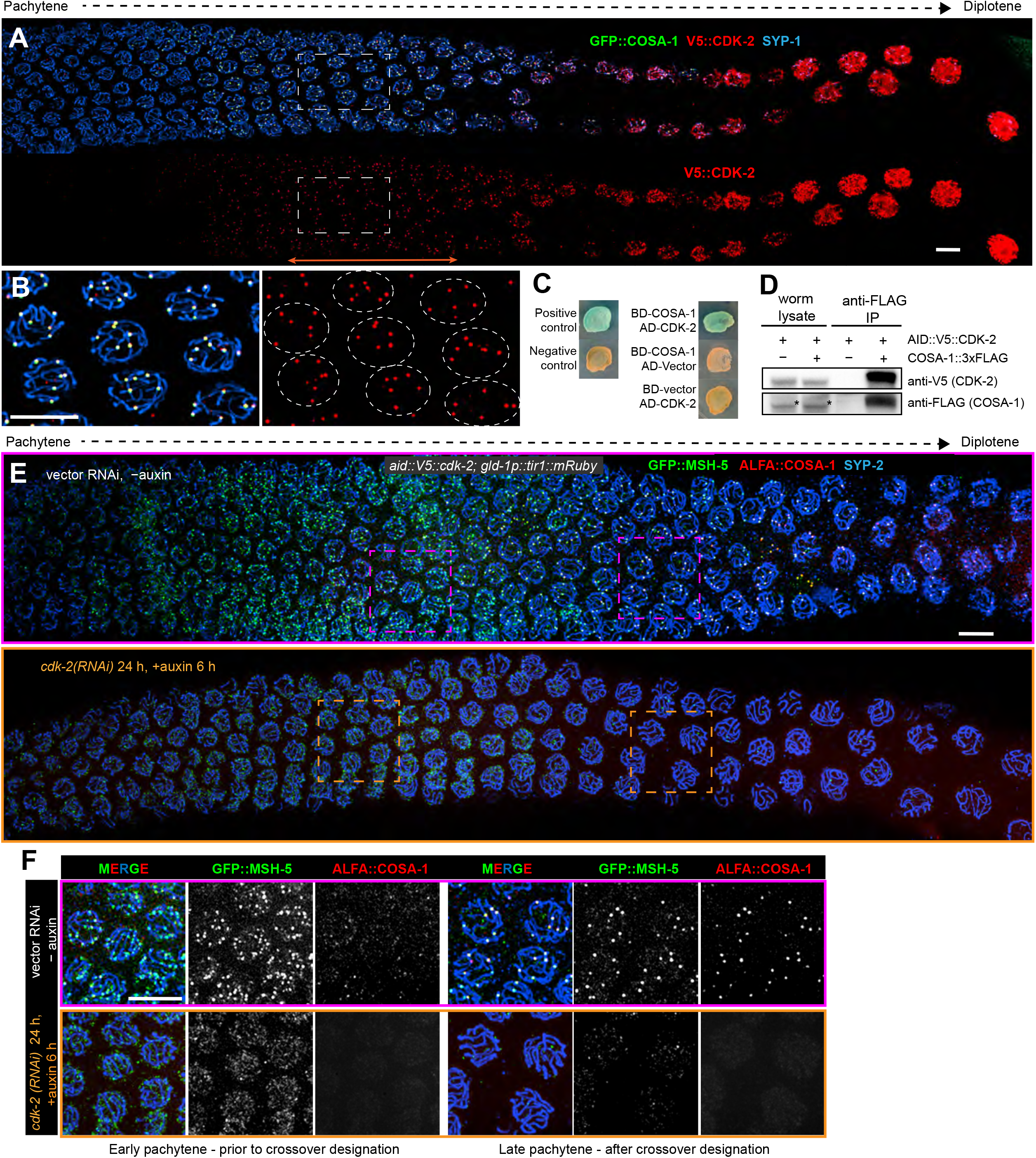
CDK-2 localizes to and stabilizes recombination intermediates. (A) Co-localization of CDK-2 with COSA-1 at designated crossover sites. The images show progression of meiotic prophase from left to right. Six foci marked by CDK-2 and COSA-1 are detected from mid-pachytene to mid-diplotene. (B) Enlarged images of the boxed regions in (A). Six foci are detected in each nucleus at late pachytene. 100% of GFP::COSA-1 foci colocalized with CDK-2 (*n*=1,017 foci from 137 nuclei and six gonads were examined). Scale bars, 5 µm. (C) CDK-2 interacts with COSA-1 in yeast two-hybrid (Y2H) assays. Here, blue patches of yeast cells indicate expression of ɑ-galactosidase resulting from physical interaction between BD-COSA-1 and AD-CDK-2 fusion proteins (see Materials and methods for details). (D) CDK-2 associates with COSA-1 in *C. elegans*. Immunoprecipitation using antibodies against the FLAG epitope strongly enriches for V5-tagged CDK-2 in lysates from a strain co-expressing COSA-1::3xFLAG, but not from a strain expressing only untagged COSA-1. COSA-1::3xFLAG is not detected in whole worm lysates due to its low abundance. Nonspecific bands recognized by the anti-FLAG antibody are indicated by asterisks. (E) Meiotic progression leads to a reduction in the number of MSH-5 foci between early and late pachytene, and appearance of six designated crossover sites marked by both MSH-5 and COSA-1 (see also Woglar and Villeneuve 2018). Depletion of CDK-2 using a combination of auxin-induced depletion and RNAi abrogates both early and late foci. Neither RNAi nor auxin-induced depletion alone was sufficient to fully deplete CDK-2 (Figure S1). Scale bars, 5 µm. (F) Enlargements of the indicated regions in (E), highlighting the larger numbers of early intermediates and six foci detected in nuclei at late pachytene, respectively. Scale bars, 5 µm.

Here we report several new experimental findings that lead us to propose a new model for crossover interference. We show that the cyclin-dependent kinase CDK-2 is required for all RNs and can be detected cytologically at designated crossover sites, along with COSA-1 and other pro-crossover factors. We find that formation of early and late foci relies on phosphorylation of the essential pro-crossover protein MSH-5, which contains many CDK consensus sites in its large, unstructured C-terminal domain. We further show that ZHP-3/4, essential pro-crossover factors conserved among all species that exhibit interference, diffuse along the SC and remain dynamic even after they accumulate within RNs. Together with other evidence, these findings indicate that recombination nodules are active biomolecular condensates whose formation, growth, and eventual spacing are regulated by posttranslational modification of component proteins. We propose a model to explain their formation and their eventual coarsening through the diffusion of proteins along the SC, resulting in wide spacing between remaining nodules. This model incorporates much of what we know about the regulation of crossover patterning across eukaryotes and naturally explains crossover interference.

## Results

### CDK-2 interacts with COSA-1 and is required for crossing-over in *C. elegans*

The discovery of COSA-1/CNTD1, a cyclin homolog, as an essential pro-crossover factor in metazoans (Holloway et al., 2014; Yokoo et al., 2012) suggested a potential role for a cyclin-dependent kinase (CDK). Cdk2 plays an essential role in crossing-over in mice (Palmer et al., 2020), implicating CDK-2 as the most likely candidate in *C. elegans*.

We detected colocalization of epitope-tagged CDK-2 with COSA-1 at foci along meiotic chromosomes from mid-pachytene through diplotene (Figures 1A and 1B). Towards the end of prophase, CDK-2 appeared much more abundant and diffusely distributed throughout the nucleoplasm, in addition to the foci detected along SCs. We also confirmed a direct physical interaction between CDK-2 and COSA-1 through co-immunoprecipitation and yeast 2-hybrid assays (Figures 1C and D).

Because CDK-2 is essential for proliferation of germline stem cells in *C. elegans*, and thus for development of the germline (Fox et al., 2011, p. 2; Jeong et al., 2011, p. 2), we used auxin-inducible degradation (AID; (Zhang et al., 2015)) to test its role during meiosis. Depletion of degron-tagged CDK-2 (AID::CDK-2) throughout the germline resulted in a dramatic reduction in crossovers, based on visualization of chromosomes at diakinesis. A few bivalents were detected at diakinesis even after 24 hours of auxin exposure, but immunostaining revealed some residual protein in germline nuclei (data not shown). To achieve more complete depletion of CDK-2, we combined *cdk-2(RNAi)*, which was insufficient to deplete CDK-2 on its own (Figure S1), with AID-mediated depletion. Following 24h of *cdk-2(RNAi)* and 6h exposure to 4 mM auxin we observed a complete loss of recombination foci marked by MSH-5 or COSA-1 (Figures 1E and 1F). ZHP-3 and ZHP-4, which normally colocalize with MSH-5 and COSA-1 at late foci, instead remained distributed along the SC (Figure 2A). Complete depletion of CDK-2 also eliminated the formation of bivalent chromosomes, indicating a failure of crossing-over (Figures 2B and 2C).

**Figure 2.**
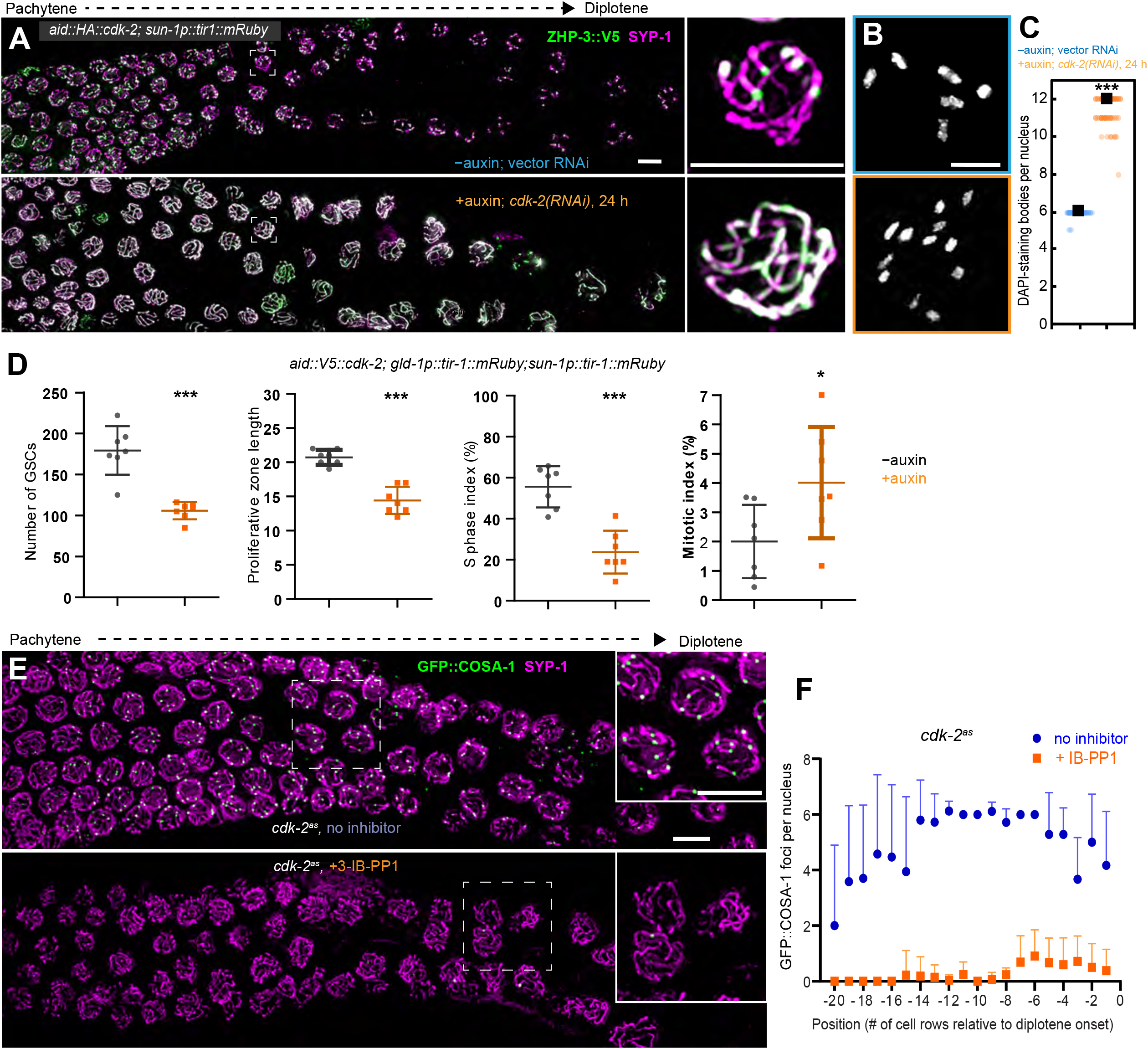
CDK-2 is required for crossing-over. (A) Images showing ZHP-3::V5 (green) and SYP-1 (magenta) in a hermaphrodite from pachytene through diplotene; regions of overlap appear white. In animals depleted of CDK-2, ZHP-3 still retained along SCs rather than concentrating at foci. Enlargements of the indicated regions are shown to the right. Scale bars, 5 µm. (B) Images of individual DAPI-stained oocyte nuclei at late diakinesis. Normally six bivalents are observed. Depletion of CDK-2 leads to an absence of chiasmata (12 univalent chromosomes). (C) Quantification of DAPI-staining bodies at diakinesis in controls and CDK-2-depleted hermaphrodites. ***p<0.0001 by Mann-Whitney test. n=56 nuclei (control) and 74 (CDK-2 depletion). Black boxes indicate median values. (D) Quantification of effects of CDK-2 depletion on germline stem cells (GSCs). Hermaphrodites homozygous for *aid::V5::cdk-2* were transferred to plates prepared with either solvent (ethanol) alone or 1 mM auxin. After 24h, animals were incubated with EdU (±auxin) and stained with antibodies that recognize histone H3 phosphorylated on serine 10 (H3pSer10) to identify cells in S-phase and mitosis, respectively (see Materials and methods). Each data point represents one gonad; data were obtained from 7 gonads for each condition. Significant reductions were observed in the total number of GSCs and the rows of GSCs observed following CDK-2 depletion; the fraction of cells in S-phase also decreased, while the mitotic index was higher in auxin-treated animals. Bars indicate mean and SD. \**p*=0.0376 and \*\*\**p*<0.0001, respectively, based on two-sided Student’s *t*-test. (E) Inhibition of CDK-2 kinase activity disrupts crossover designation. Hermaphrodites homozygous for an analog-sensitive *cdk-2* allele (*cdk-2^as^*) were treated with the ATP analog 3-IB-PP1 for 24 hours. Immunostaining shows strongly reduced numbers of GFP::COSA-1 foci along SCs (SYP-1 immunofluorescence, magenta). Enlarged images of the indicated regions are shown as insets. Scale bars, 5 µm. (F) Quantification of GFP::COSA-1 foci in late pachytene nuclei in *cdk-2^as^* hermaphrodites exposed to 8 µg/ml 3-IB-PP1 for 24 hours. The X-axis shows nuclear position as the number of cell rows relative to diplotene onset. GFP-COSA-1 foci per nucleus is plotted as mean+SD values.. Data were collected from 6 and 12 gonads for control and treated animals (*n* = 243 and 420 nuclei, respectively).

Germline stem cells (GSCs) in gonads depleted of CDK-2 showed severe defects in proliferation (Figure 2D), consistent with the role of CDK-2 in GSC mitosis (Fox et al., 2011, p. 2; Jeong et al., 2011, p. 2). Crossovers in *C. elegans* depend on homolog pairing and synapsis (Colaiácovo et al., 2003; MacQueen, 2002), both of which occurred normally in the absence of CDK-2 (Figures S2B and S2C). CDK-2 depletion also resulted in an extended region of meiotic nuclei containing RAD-51 foci, indicating that programmed DSBs do not depend on CDK-2 (Figures S2D and S2E). A similar extension of RAD-51-positive foci is observed in a wide variety of mutants with defects in crossover recombination; a failure to establish crossover intermediates activates a crossover assurance checkpoint that prolongs DSB induction (Yu et al., 2016). Together this indicates that CDK-2 is dispensable for the checkpoint, and is specifically required to promote or stabilize the association of pro-crossover factors with recombination intermediates.

We also constructed an analog-sensitive allele (Lopez et al., 2014), *cdk-2^as^*, to investigate whether its kinase activity is required for recombination nodules (see Materials and methods). Exposure of control worms to the ATP analog 3-IB-PP1 did not cause any meiotic defects, but the same treatment resulted in greatly reduced numbers of LRNs and chiasmata in *cdk-2^as^* homozygotes (Figures 2E and 2F; and data not shown). Thus, the kinase activity of CDK-2 is essential for crossing-over.

### Crossing-over does not depend on most canonical CDK-2 regulatory pathways

The mitotic activity of Cdk2/CDK-2 is regulated by several mechanisms that together mediate switch-like activation of the kinase and coordinate its activity with cell growth (Morgan, 1995). Since many of the factors involved with this regulation are essential for development and germline proliferation, we tested whether they contribute to crossing-over or the appearance of recombination foci by tagging the endogenous genes with the AID degron and depleting the proteins in the germline. In most cases, effective depletion could be validated by examining the effects on proliferating GSCs in the distal region of the germline.

While the meiosis-specific cyclin homolog COSA-1 is essential for crossovers (Yokoo et al., 2012), we found that other cyclins that can regulate CDK-2 do not contribute to crossover formation. Depletion of cyclin A (CYA-1) or cyclin E (CYE-1) in the germline did not impair meiotic progression, crossover designation, or bivalent formation (Figures S3A-S3C and S3E-S3G). Depletion of CYE-1 resulted in dramatic reduction of GSCs in S-phase (Figure S3D), consistent with the established role of CDK-2/CYE-1 in driving the G1/S transition in GSCs (Fox et al., 2011; Jeong et al., 2011). Depletion of CYA-1 did not result in any apparent germline defects (Figures S3E-S3H).

Full activation of CDK-2/cyclin complexes usually requires phosphorylation of the activation loop of CDK-2 by a CDK-activating kinase (CAK) and removal of inhibitory phosphorylation near Cdk2 N-terminus by the Cdc25 phosphatase. We depleted CDK-7, the core catalytic component of CAK (Harper and Elledge, 1998), in the germline (Figure S4A). Surprisingly, CDK-7 depletion caused a more severe cell cycle arrest among GSCs than depletion of CDK-2, based on the number of cells detected after 24 hours (Figures S4B-S4D). Depletion of CDK-7 also disrupted meiotic progression, as indicated by some synapsis defects and a block to pachytene exit (Figure S5). This may reflect a role for CDK-7 in activating CDK-1 or other kinases. Nevertheless, crossover designation still occurred following depletion of CDK-7; six COSA-1 foci were detected in many late prophase nuclei, although the spatial distribution of nuclei displaying these foci was somewhat perturbed (Figures S5 and S6).

We also tested whether crossover designation or chiasma formation might be regulated by phosphorylation or dephosphorylation of CDK-2 at conserved N-terminal residues (T25/Y26) that control its mitotic activity. We tagged all four genes encoding *C. elegans* homologs of Cdc25 phosphatase (*cdc-25.1, cdc-15.2, cdc-25.3* and *cdc-25.4*) with the AID degron. Co-depletion of these 4 paralogs resulted in severe mitotic defects, including an absence of S-phase nuclei and the presence of a few large, likely polyploid GSCs (Figures S7A-S7D). However, crossover designation was unimpaired (Figures S7E and S7F). Similarly, mutation of these sites to nonphosphorylatable residues (T25A; Y26F) did not affect crossover designation (Figures S7G and S7H).

Taken together, we conclude that COSA-1 is probably the only cyclin required for the essential meiotic activity of CDK-2 in *C. elegans*. This differs from meiosis in mammals, where Cdk2 plays important roles at telomeres prior to crossover designation, for which it requires a different cyclin-like protein, RINGO/Speedy (Tu et al., 2017; Viera et al., 2014). However, the COSA-1 ortholog CNTD1 is essential for crossing-over in mammals and like COSA-1, colocalizes with Cdk2 at recombination sites (Holloway et al., 2014). COSA-1 may promote activation of CDK-2 without phosphorylation of its activation loop, as has been described for Ringo/SPEEDY (McGrath et al., 2017), or CDK-2 activation may require phosphorylation by a kinase other than CDK-7.

### Phosphorylation of MSH-5 is required to stabilize recombination intermediates

As described in the Introduction, pro-crossover factors can be detected at early and late foci in *C. elegans* GFP::MSH-5 (Janisiw et al., 2018) is currently the best marker for cytological detection of early crossover intermediates, while epitope-tagged COSA-1, CDK-2, ZHP-3, and ZHP-4 can all be detected at the late site (Woglar and Villeneuve, 2018). Early and late foci also depend on ZHP-3/4 (Zhang et al., 2018) and CDK-2 (above), suggesting that interactions among all these pro-crossover proteins drive the formation of both early and late sites. We considered how CDK-2 might promote such interactions. CDK substrates typically contain clusters of potential phosphorylation sites (S/T-P) (Moses et al., 2007). MSH-5 in *C. elegans* and other *Caenorhabditids* has a very long C-terminal extension that is predicted to be disordered (Figure S8). This domain contains between 9-24 S/T-P motifs (predominantly T-P) in different species, with 13 in *C. elegans* MSH-5. MSH-5 can also be phosphorylated in *vitro* by human CDK1 (Yokoo et al., 2012) and CDK2 (data not shown) and was thus a prime candidate for CDK-2 regulation.

We tested the potential effects of MSH-5 phosphorylation by editing the endogenous gene using CRISPR/Cas9. Because we expected that phosphorylation of these sites might have additive effects, we designed templates to replace blocks of coding sequence containing several candidate sites with nonphosphorylatable or phosphomimetic residues (Figure 3A). “Block 1” includes five candidate sites (T1009, S1029, T1046, T1057, and T1094), while “Block 2” includes six nonoverlapping sites (T1109, T1126, T1164, T1253, S1278, and T1295). Starting with a strain in which the endogenous *msh-5* locus was tagged at its N-terminus with the GFP coding sequence (Janisiw et al., 2018), we generated four mutant alleles by mutating each block of residues to either alanine (nonphosphorylatable, NP) or aspartate (phosphomimetic, PM). We designated the mutant proteins as MSH-5^NP1^ and MSH-5^NP2^ (nonphosphorylatable substitutions in Block 1 or Block 2, respectively), and MSH-5^PM1^ and MSH-5^PM2^ (phosphomimetic mutations at the corresponding sites). We confirmed germline expression of mutant proteins by imaging intact, living animals (Figure 3B). We analyzed the effects of these mutations on protein localization and function (Figures 3 and 4).

**Figure 3.**
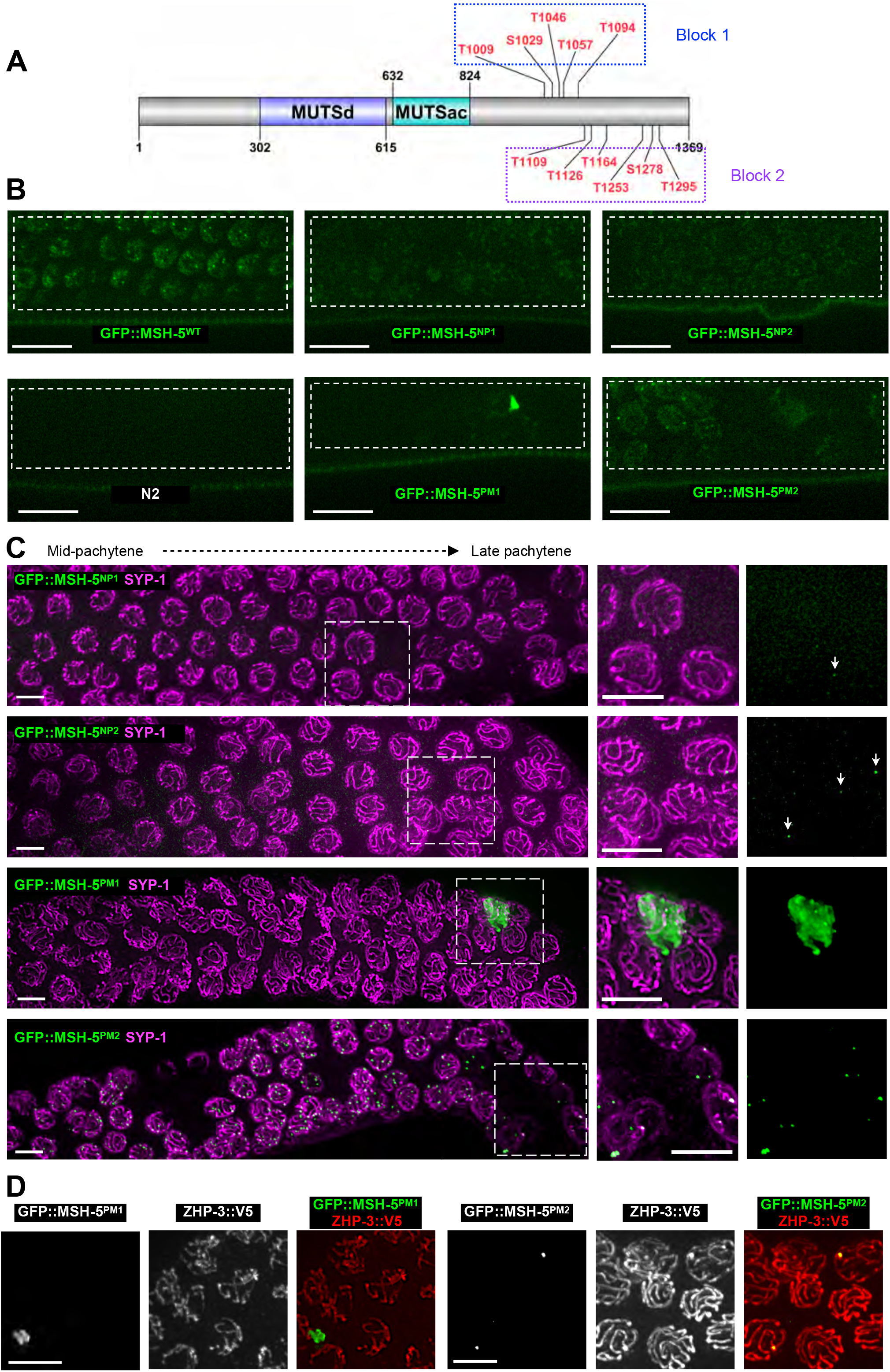
Mutation of putative phosphorylation sites in MSH-5 disrupts protein localization. (A) Diagram of MSH-5 showing the DNA binding (MUTSd) and ATPase (MUTSac) domains. The positions of 11 potential CDK sites that were mutated to alanine (nonphosphorylatable; NP) or aspartate (phosphomimetic, PM) in the unstructured C-terminal domain are shown. “Block 1” includes 5 sites that were mutated in MSH-5^NP1^ and MSH-5^PM1^; “Block 2” sites were mutated in MSH-5^NP2^ and MSH-5^PM2^. (B) Confocal images of the pachytene region of germlines in live animals expressing GFP::MSH-5 (wild-type) and each of the phosphosite mutants. GFP-labeled protein is detected in all mutants. Its distribution is more diffuse in the two alleles with nonphosphorylatable substitutions, and aberrant foci are seen in the two phosphomimetic alleles. N2 (wild-type, no GFP::MSH-5 expression) is shown as a negative control. Images were collected and scaled identically. (C) Immunofluorescence of GFP::MSH-5 and SYP-1 (SC) in germlines from each of the phosphosite mutants, with the indicated regions enlarged on the right. Nonphosphorylatable substitutions in MSH-5 result in severe reduction in the appearance of foci along chromosomes; a few remaining foci associated with SC are highlighted and indicated with arrowheads in the enlarged views. The MSH-5^PM1^ mutant protein usually forms a single large aggregate in each animal, usually in extracellular space within the germline. MSH-5^PM2^ localizes to foci along SCs in mid-pachytene, but these decrease in number by late pachytene. Ectopic foci not associated with SCs or nuclei are also detected in this mutant. Scale bars, 5µm. (D) Localization of ZHP-3::V5 in *msh-5* phosphomimetic mutants. ZHP-3 does not colocalize with the large aggregates of GFP::MSH^PM1^. It does concentrate at foci formed by GFP::MSH^PM2^ along SCs, but it also remains distributed along the SC in this mutant, unlike in wild-type meiosis. Scale bars, 5 µm

**Figure 4.**
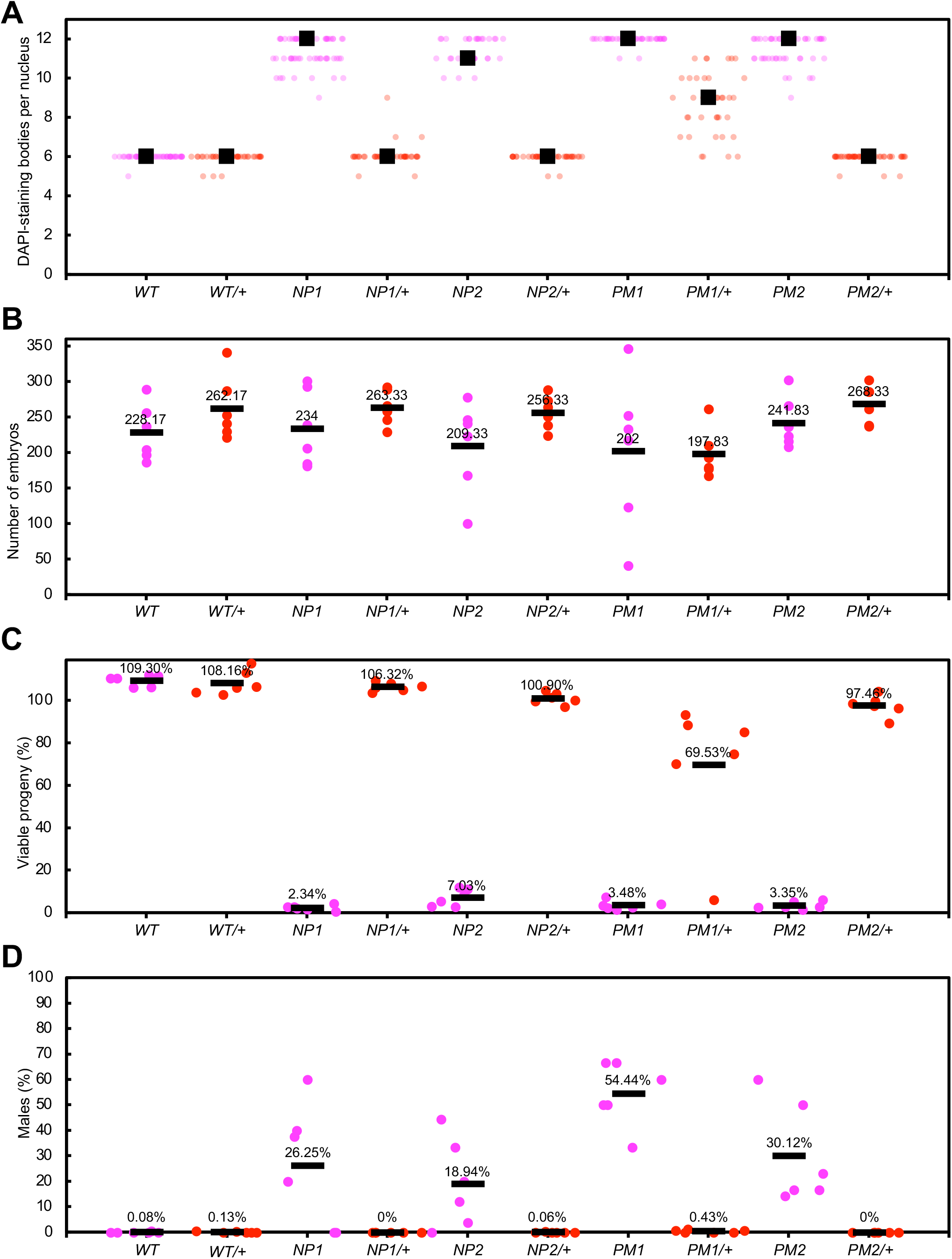
Quantification of chiasmata and broods produced by *msh-5* phosphosite mutants. (A-D) All graphs show data for homozygous and heterozygous animals expressing wild-type GFP::MSH-5 (*WT*) and the 4 mutant alleles (*NP1, NP2, PM1,* and *PM2*). In all heterozygotes, the “+” chromosome was the *tmC9* balancer. (A) Chiasma formation was assessed by quantification of DAPI-staining bodies at diakinesis. Normal crossing-over results in the appearance of 6 bivalents (DAPI-staining bodies), while crossover defects result in up to 12 univalent chromosomes. Each data point (dot) represents one nucleus; median values are indicated by boxes. (B-D) Brood analysis. Each data point represents one full brood; six broods were analyzed for each genotype. Mean values are indicated by the bars and numbers. (B) Brood sizes (number of embryos). (C) Viable progeny, expressed as a ratio of L4/adult progeny over total embryos laid. Values exceeding 100% are common because embryos are small and some escape detection. (D) Male self-progeny expressed as a ratio of males to all viable offspring. Males have a single X chromosome and arise among hermaphrodite self-progeny due to X nondisjunction.

The phosphosite substitutions caused dramatic effects on the distribution of MSH-5 protein (Figures 3B and 3C). Both mutants with nonphosphorylatable substitutions showed greatly reduced localization of MSH-5 to foci along meiotic chromosomes, although diffuse nucleoplasmic fluorescence was detected. In contrast, both proteins with phosphomimetic substitutions formed aberrant aggregates. MSH-5^PM2^ localized to some foci along SCs. These were irregular in number and gradually declined to ∼1 per nucleus by diakinesis. We also observed occasional ectopic foci that were not associated with SCs or chromosomes. Animals expressing MSH-5^PM1^ showed even more dramatic mislocalization: the mutant protein formed very large, stable aggregates, often several microns in diameter, in extracellular regions of the germline.

Hermaphrodites homozygous for each of the four mutant alleles showed severe defects in meiotic chromosome segregation, based on their production of many inviable embryos and a high incidence of male self-progeny, a.k.a. the “Him” phenotype 8/26/21 7:17:00 PM (Figure 4). Most produced only a few more viable progeny than the 2.5% viability reported for *msh-5* null mutants (Figure 4; (Kelly et al., 2000)). Both alleles containing nonphosphorylatable substitutions behaved recessively. In contrast, expression of MSH-5^PM1^ caused some chromosome missegregation even in heterozygotes, and a weak dominant effect was also detected for MSH-5^PM2^.

In *msh-5^pm1^* mutants, ZHP-3 remained distributed throughout the SC rather than colocalizing with the large MSH-5 aggregates at late prophase (Figure 3D). ZHP-3 did concentrate with MSH-5^PM2^ at some foci associated with SC, but also remained throughout the SC. The absence of chiasmata in these mutants (Figure 4), indicates that the aggregates were insufficient to promote crossover resolution.

We interpret these findings to indicate that phosphorylation of MSH-5 promotes aggregation and perhaps interactions with other pro-crossover factors, and that reduced phosphorylation prevents accumulation of pro-crossover factors, leading to noncrossover resolution of all recombination intermediates.

### Protein dynamics at recombination sites and along SCs

Our previous work showed that the RING finger proteins ZHP-3 and ZHP-4 interact with each other and show dynamic localization during prophase, first along the SC and later concentrated at designated crossover sites (Zhang et al., 2018). Evidence that the synaptonemal complex (SC) behaves as a liquid crystalline material (Rog et al., 2017) also suggested that proteins might diffuse along this compartment. Previous studies have demonstrated the dynamic localization SC proteins using photoconversion and photobleaching assays, but these have limited ability to quantify the directionality or rate of movement, particularly for highly dynamic proteins.

To investigate the dynamics of pro-crossover proteins within RNs, we used fluorescence recovery after photobleaching (FRAP). We selectively bleached individual late nodules (Figure 5A) and analyzed their fluorescence recovery in worm strains expressing fluorescently tagged pro-crossover proteins. This revealed that proteins within late nodules have widely varying dynamics (Figure 5B). CDK-2, COSA-1 and MSH-5 were all relatively static, as evidenced by an absence of fluorescence recovery. In contrast, ZHP-3 and ZHP-4 showed rapid recovery, indicating that these proteins remain mobile even after their accumulation at LRNs, and likely exchange with proteins along the SC. Together with evidence that the formation of RNs is regulated by phosphorylation (above), these findings suggest that recombination nodules are active biological condensates that contain both highly dynamic and more stable components.

**Figure 5.**
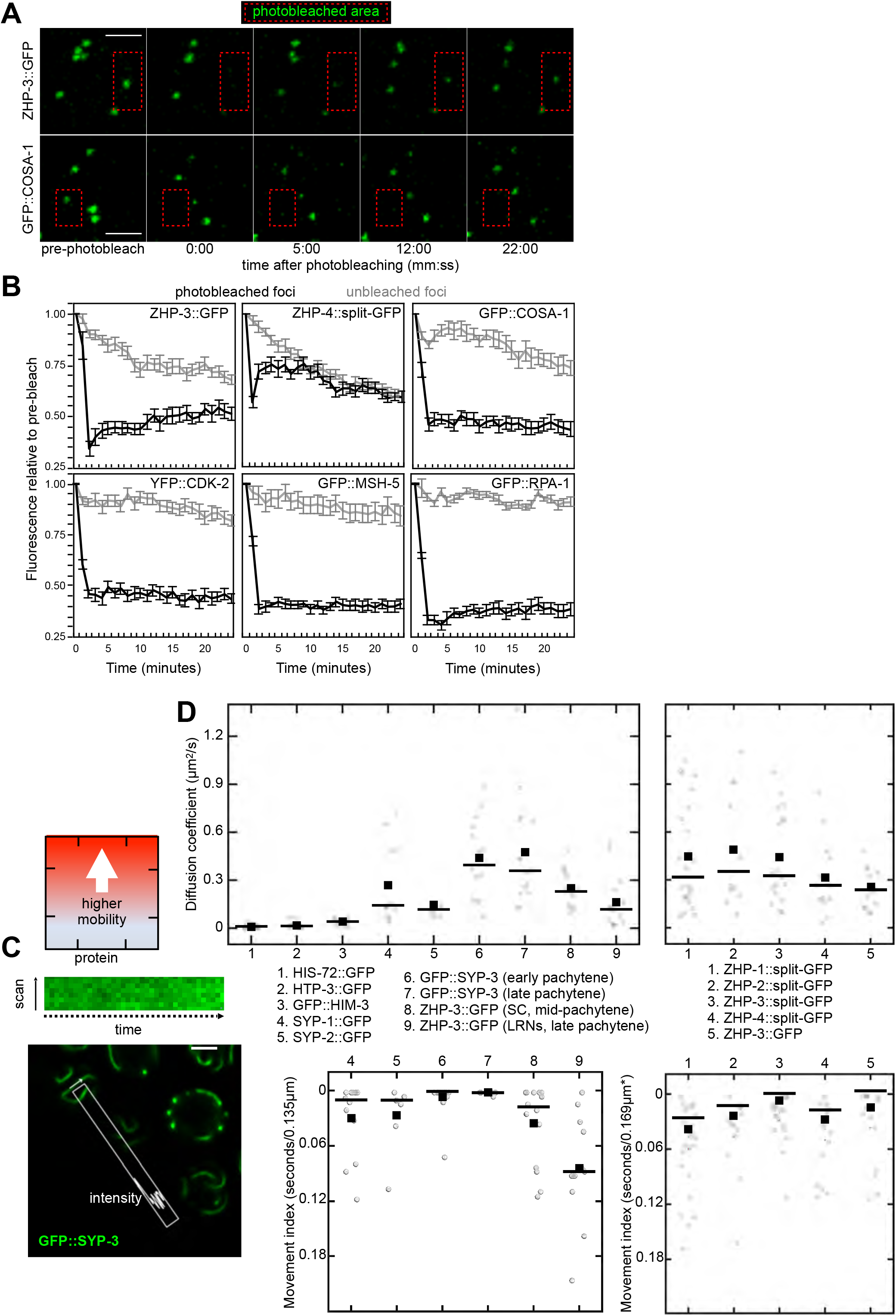
Protein dynamics at designated crossover sites and along SCs. (A)-(B) Analysis of protein dynamics at recombination sites using fluorescence recovery after photobleaching (FRAP) (A) Designated crossover foci in living worms expressing fluorescently-tagged proteins were selected for photobleaching. Projection images for ZHP-3::GFP and GFP::COSA-1 are shown as examples. ZHP-3::GFP intensity recovers after photobleaching fluorescence recovery, while GFP::COSA-1 shows little or no recovery. Scale bar, 2µm. (B) Quantification of fluorescence after photobleaching. Integrated intensities for each focus were normalized against the intensity of the same region before bleaching. Unbleached foci from the same nuclei were also quantified; these underwent slower photobleaching due to time-lapse image acquisition. Averages were calculated for both bleached and unbleached foci. Error bars show the standard error of the mean. Numbers of foci analyzed for each protein were as follows: ZHP-3::GFP (17 bleached, 18 unbleached), ZHP-4::split-GFP (14 bleached, 15 unbleached) GFP::COSA-1 (21 bleached, 19 unbleached), YFP::CDK-2 (18 bleached, 20 unbleached), GFP::MSH-5 (20 bleached, 19 unbleached). GFP::RPA-1 (19 bleached, 20 unbleached). (C) Illustration of our data collection approach. For data shown in (D) (also in Figure S9), higher values indicate greater protein mobility/faster diffusion. Data were collected as time series of short (1 µm) point scans along individual SCs. See Materials and methods for details. (D) Labels indicate the fluorescent proteins analyzed and apply to the graphs above (showing diffusion constants) and below (showing movement indices). Each data point is an estimated diffusion coefficient (above) or movement index (below) based on analysis of one data set. All diffusion coefficients were based on data from nuclei in late pachytene, unless otherwise noted. Mean and median values are indicated by bar boxes and bars, respectively. Data for ZHP-1-4 were obtained using a different microscope and at an earlier stage of meiosis (prior to crossover designation) and are thus plotted separately. The same data for ZHP-3::GFP are plotted twice to facilitate comparison with ZHP-3::split-GFP. Movement indices could not be calculated for the first three proteins because they move too slowly.

To understand the movement of proteins along the SC, we developed an approach based on fluorescence correlation spectroscopy (FCS) (Hinde et al., 2011). Living worms expressing fluorescently tagged proteins were immobilized and imaged using a point-scanning confocal microscope (Figure 5C). We identified regions of SC lying parallel to the plane of the coverslip and acquired 1-dimensional scans along short (1-µm) segments of these SCs at high speed (473 µs/scan at 2114 Hz). The fluctuations observed over time in these scans were analyzed to calculate their autocorrelation functions (ACFs) and paired correlation functions (pCFs), which enabled estimates of diffusion rates and motion of proteins within the SC, respectively (Figures 5D, 6, and S9).

**Figure 6.**
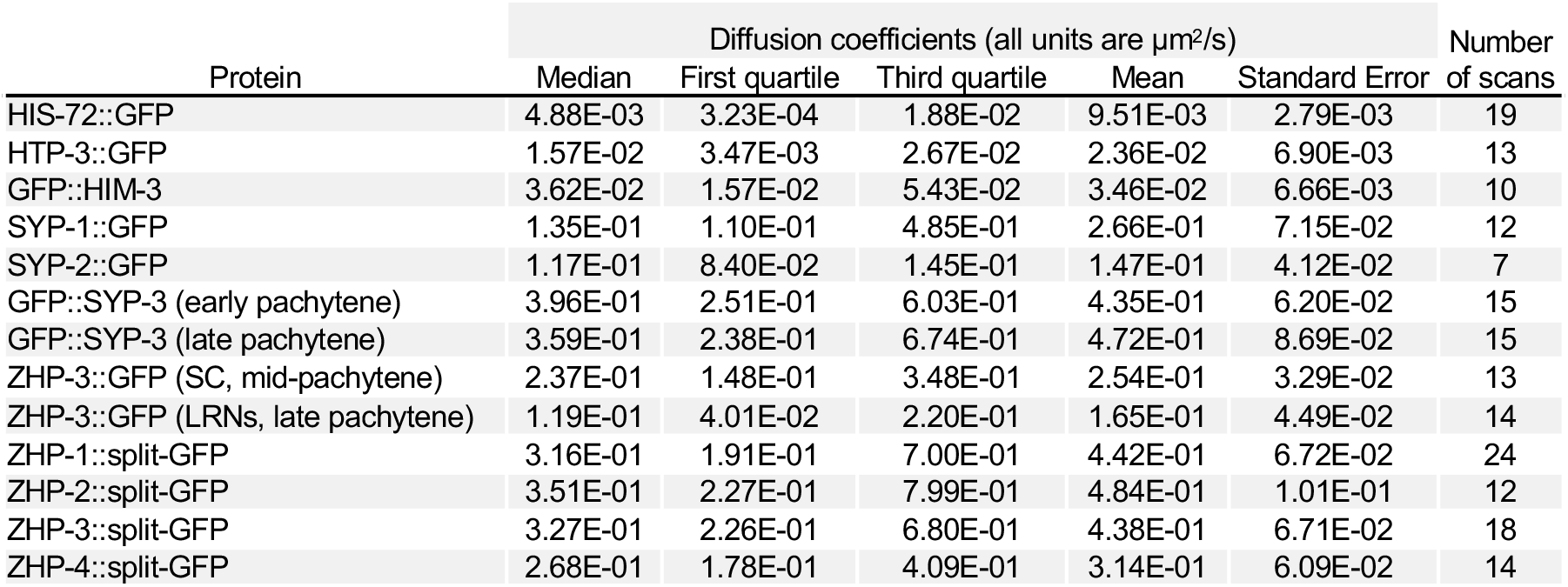
Summary of diffusion coefficients measured by FCS. A table containing useful summary values for all diffusion coefficients in Figure 5D.

We first validated the method by estimating diffusion coefficients for relatively static proteins whose movement has been previously analyzed. These included a histone, H3.3 (GFP::HIS-72) and two HORMA proteins associated with meiotic chromosome axes, HTP-3 and HIM-3 (Figure 5D). HIS-72::GFP showed a median estimated diffusion coefficient of 4.88×10^−3^ µm^2^/s (Figure 6), consistent with other measurements of chromatin mobility. Our estimated diffusion coefficients for HTP-3 and HIM-3 were somewhat higher, 1.57×10^−2^ µm^2^/s and 3.62×10^−2^ µm^2^/s, respectively. These values are consistent with measurements based on single-particle tracking of HTP-3 (von Diezmann and Rog, 2021) and previous evidence that HIM-3 is less dynamic than SYP-3 (Rog et al., 2017). The slightly higher mobility of HIM-3 compared to HTP-3 is also consistent with evidence that HIM-3 is recruited to the axis by binding HTP-3, while HTP-3 likely binds directly to cohesins that form the axis core (Kim et al., 2014).

We next measured dynamics for three of the six known SC proteins, starting with SYP-3. Two studies that used FRAP to analyze SYP-3 mobility within the SC reported that GFP::SYP-3 recovers to a greater extent in early pachytene than in late pachytene (Nadarajan et al., 2017; Pattabiraman et al., 2017). This could reflect either reduced mobility of SYP-3 within the SC or a more limited pool of free protein in the nucleoplasm that can exchange with proteins in the SC. We analyzed SYP-3 diffusion in both early and late pachytene nuclei and found no significant differences (Figures 5D, S9, and Table S5), supporting the latter interpretation. Consistent with this, the fluorescence intensity of SYP-3 within the SC increases more than 2-fold between early and late pachytene (Pattabiraman et al., 2017) while its nucleoplasmic fluorescence decreases (not shown). Two other SC proteins, SYP-1 and SYP-2 also showed higher mobility than axis proteins, albeit somewhat lower than SYP-3 (Figures 5D and 6).

We also measured diffusion of the four RING domain proteins required for crossover patterning, ZHP-1-4. All four proteins localize throughout SCs from early prophase until mid-pachytene, when ZHP-3/4 begin to concentrate at designated crossover sites (Zhang et al., 2018). We found that the mobility of all four proteins along the SC was similar to those of SYP-1 and SYP-2. ZHP-3 showed somewhat lower mobility after concentrating at recombination foci, but remained dynamic, consistent with our FRAP analysis, above. All of the diffusion coefficients derived from this analysis are summarized in Figure 6.

In addition to autocorrelation function analysis of our line-scan data, we calculated the paired correlation functions (pCFs) of fluorescence measured between voxels. This method determines the average time required for fluorescent proteins to move between voxels at a given distance, which correspond to 0.135 µm or 0.169 µm in our data sets (see Materials and methods). This approach is well-suited to measuring directional movement of proteins along the SC or within RNs. It is complementary to and independent of ACF calculations, although it is applied to the same data (Digman and Gratton, 2009).

We calculated the average pCF for each scan (Figure 5D), as well as for each pair of adjacent pixels in the datasets (Figure S9B). The measured values were highly concordant with the diffusion coefficients we measured through ACF analysis, indicating that most of these proteins’ diffusion occurs along the SC, although we cannot rule out some exchange with the nucleoplasm. These measurements also corroborated our conclusion that the mobility of these proteins along the SC does not change significantly between early and late pachytene.

### An active droplet model for the formation of recombination nodules

Cytological studies of recombination nodules have suggested that they may be biomolecular condensates. In organisms where they have been observed by electron microscopy, both early and late nodules appear as ellipsoidal or spherical bodies associated with, but not embedded within, SCs (Carpenter, 1975; Schmekel and Daneholt, 1998; Wettstein et al., 1984; Zickler, 1977). We have also found that ZHP-3/4 remain dynamic within recombination foci. Our evidence that CDK-2 is essential for RN formation, and that a phosphomimetic form of MSH-5 spontaneously forms aggregates even outside of cells, lead us to propose an active droplet model of RN formation and dynamics (Figure 7A) In this model, RN formation requires phosphorylation of at least one component – MSH-5 in *C. elegans*. MSH-5 is initially recruited to JMs associated with axes or SCs through the DNA binding activity of MutSɣ (HIM-14/MSH-5 in *C. elegans*). Phosphorylation by CDK-2 stabilizes the protein at these sites and promotes its activity as a scaffold to recruit additional RN proteins, which partition into nodules as clients (Banani et al., 2016; Wheeler and Hyman, 2018). Crucially, only highly phosphorylated MSH-5 is stably associated with recombination sites, so that controlling its phosphorylation regulates RN formation (Kirschbaum and Zwicker, 2021; Söding et al., 2020).

**Figure 7.**
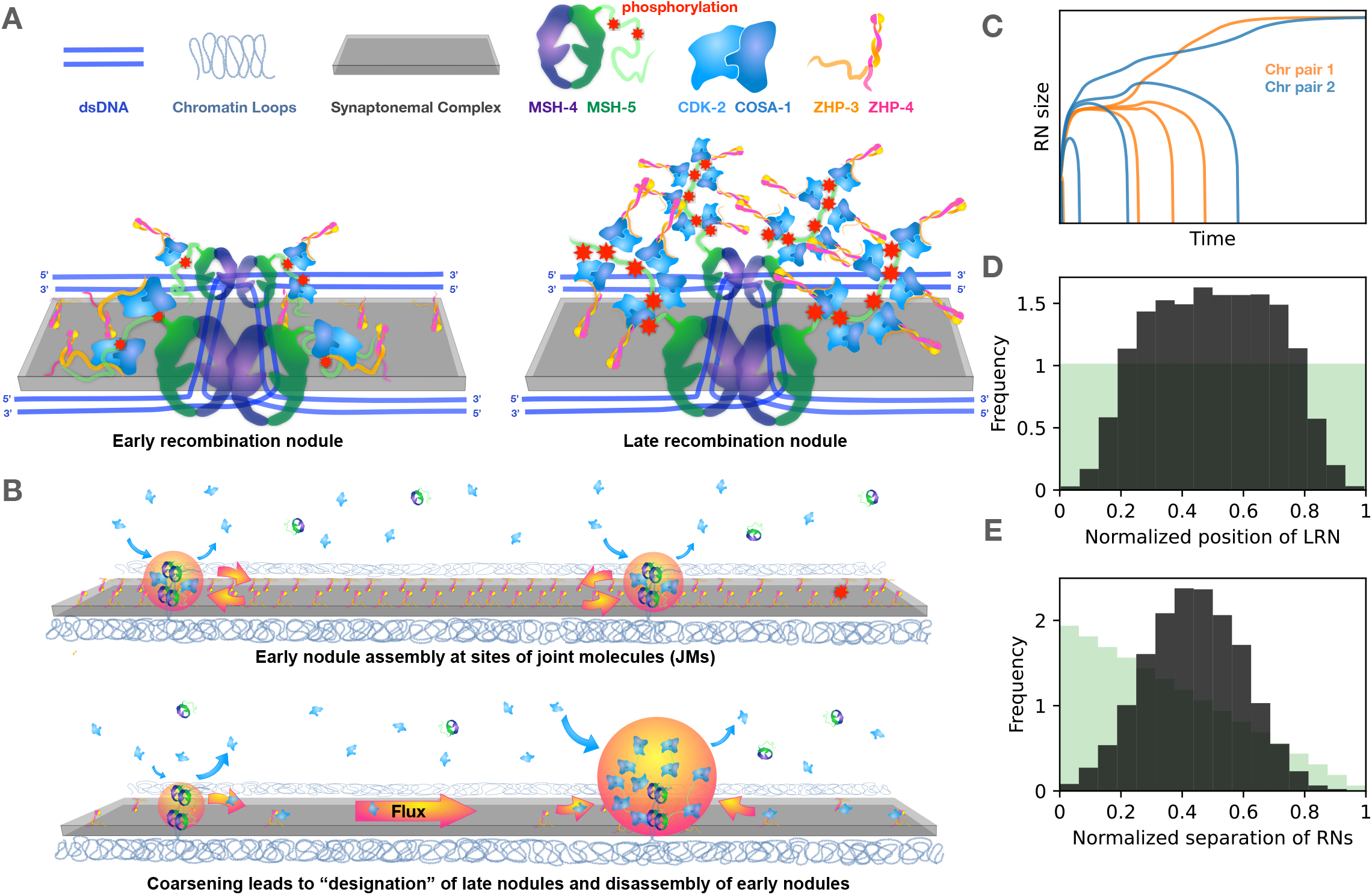
Model for organization and coarsening of recombination nodules. (A) A speculative schema for the organization of early and late recombination nodules. JMs are illustrated as double Holliday Junctions, based on molecular evidence primarily from *S. cerevisiae*, and on cytological evidence from *C. elegans* that recombination sites have two-fold symmetry, but both this structure and its physical disposition relative to the SC are speculative. Only the two recombinant chromatids that are connected by the JM are illustrated, and chromatin loops flanking the SC are omitted for simplicity. Protein shapes are based loosely on known structures and/or AlphaFold predictions (Jumper et al., 2021). MSH-5, ZHP-3, and ZHP-4 all have C-terminal domains that are predicted to be disordered. MSH-5 is phosphorylated on its C-terminal tail by CDK-2, which promotes nodule stability and grow through recruitment of ZHP-3/4 and additional CDK-2/COSA-1. A direct physical interaction between CDK-2/COSA-1 and MSH-4/5 is entirely speculative, but their depiction within the nodule “core” is suggested by our FRAP evidence that these components are relatively stable within nodules, while ZHP-3/4 continually exchange between nodules and the SC. Physical interaction between ZHP-3/4 and CDK-2/COSA-1 is also hypothetical but illustrates our evidence that ZHP-3/4 are for CDK-2 activity, and that ZHP-3/4 and CDK-2/COSA-1 both accumulate as nodules coarsen, while the fluorescence intensity of MSH-5 is similar between early and late nodules. (B) Illustration of our model for the initial formation and coarsening of recombination nodules. The synaptonemal complex (SC) is shown as a gray slab that acts as a conduit for diffusion of ZHP-3/4 along the interface between homologous chromosomes, which are depicted as chromatin loops. Assembly of early nodules is initiated by binding of MSH-4/5 to joint molecules (JMs) that are arise through repair of double-strand breaks. CDK-2/COSA-1 are recruited and activated by a mechanism that requires ZHP-3/4, which are associated with the SC. CDK-2 phosphorylates MSH-5 within its C-terminal tail, which stabilizes the early nodules. Nodules reach a steady state if protein loss due to diffusion and/or dephosphorylation equals the rate of recruitment. Coarsening of nodules arises through competition for proteins that exchange via the SC. Recruitment of active CDK-2 creates a feedback loop promoting further recruitment of CDK-2/COSA-1 and ZHP-3/4. Larger nodules recruit proteins more rapidly, leading to depletion of proteins from adjacent nodules until only one LRN remains along each SC. Dissolution of nodules leads to noncrossover resolution of the JMs at those sites, while persistence of pro-CO proteins at the late nodules leads eventually to recruitment of resolvases and CO resolution. (C) Sizes of RNs on two chromosome pairs as a function of time based on simulation of coarsening. All RNs start the process at a similar size, but only one RN survives per chromosome. (D) Histogram of the position of the remaining RN normalized to the length of the SC, assuming a homogeneous distribution prior to coarsening. Nodules near SC termini are more easily lost due to the lower availability of limiting components near the ends. (E) Histogram of distances between the last two remaining RNs on the same chromosome pair normalized to the length of the respective SC. Ten RNs were placed uniformly along the SC and the distance between the last two RNs was recorded for 10^5^ independent repetitions. Green bars indicate the expected distribution of distances between two randomly placed RNs.

Formation of all RNs also requires ZHP-3/4, which are confined to the SC (Zhang et al., 2018). We thus propose that ZHP-3/4, along with COSA-1, are required to activate CDK-2. Recruitment of the MutSƔ complex to JMs is accompanied by phosphorylation of MSH-5, thereby stabilizing its localization and promoting formation of early recombination nodules. Phosphatase activity in the nucleoplasm ensures that MSH-5 does not spontaneously self-assemble. Binding of MutSƔ to JMs and their association with the SC, together with a scaffolding activity of MSH-5, is sufficient to explain the condensation of proteins to form ERNs at sites of JMs.

These aspects of the model account for the evidence that RNs only form in the presence of MutSɣ, CDK-2, COSA-1, and ZHP-3/4. Additionally, they explain why RN formation is impaired when phosphorylation sites on MSH-5 are mutated to nonphosphorylatable residues, while MSH-5 accumulation cannot be spatially controlled if it is constitutively phosphorylated. The model further predicts that the size of ERNs is dictated by the balance of material influx (proportional to the local rate of MSH-5 phosphorylation) and material efflux (either by spontaneous unbinding or by dephosphorylation in the ERN).

The requirement for a localized kinase activity enables ERNs to coexist stably at multiple sites along the same SC. While in principle RNs could form through spontaneous phase separation at sites that recruit a scaffold protein, their sizes would typically be unstable in such a passive system. This instability arises due to surface (interfacial) tension of phase-separated bodies, which favors the transfer of components from smaller to larger condensates (Hyman et al., 2014). The resulting coarsening process is known as Ostwald ripening. This well-studied behavior of multiphase systems can be inhibited if condensation depends on modification by a localized activity, since the direct exchange of material between condensates is then suppressed (Söding et al., 2020; Weber et al., 2019; Zwicker et al., 2015). However, if this activity were able to exchange via the nucleoplasm or SC, this would also lead to RN instability. We thus propose that active CDK-2 is initially restricted in its ability to move between recombination nodules, thereby ensuring that ERNs form independently of each other.

### Coarsening of recombination nodules can explain crossover interference

We next extend our model of RN formation to explain the process leading to crossover designation and interference. A conserved phenomenon that the model must explain is that only a few of the ERNs eventually become LRNs; in *C. elegans* this is usually limited to one per chromosome pair. While the switch between ERNs and LRNs is governed by a nucleus-wide signal, the selection of LRNs happens for each chromosome pair independently. This implies that ERNs on the same chromosome pair communicate with each other, while they act independently of ERNs on other chromosome pairs.

These features can be explained if CDK-2 activity moves along SCs and accumulates in nodules, driving further growth. Through the ripening process described above, this would lead most RNs to lose CDK-2 activity and shrink while some RNs gain activity, grow, and become LRNs (Figures 7B and 7C). If CDK-2 activity were exchanged through the nucleoplasm, such a ripening process would eventually lead to a single surviving RN per nucleus, rather than one per SC. However, if diffusion occurs only along SCs, RNs can only compete with other RNs along the same SC. Competition is reduced as the distance between nodules increases due to the time required for diffusion and the presence of intervening nodules. These aspects of the model are similar to the diffusion-mediated coarsening of Hei10 that was recently proposed for *Arabidopsis* (Morgan et al., 2021). These models predict that RNs at the end of the SC are less likely to survive (Figure 7D) and that surviving LRNs are spaced farther apart than randomly selected ERNs (Figure 7E), *i.e*., that they show interference. Multiple LRNs may remain along the same SC if crossover resolution takes place before coarsening reaches completion. Stability of multiple LRNs per SC could also arise if there is an upper bound to the amount of material that can accumulate within each RN. Taken together, our model accounts for patterning of LRNs through a coarsening process, which is mediated by the transfer of CDK-2 activity between nodules along the SC.

## Discussion

### The role of the SC in crossover interference

A variety of physical and mathematical models have been proposed to explain crossover interference (reviewed by Otto and Payseur, 2019; Pazhayam et al., 2021). An early model suggested that the SC might behave as a “compartment” within which recombination proteins might diffuse and accumulate cooperatively at a limited number of recombination sites (Holliday, Robin, 1977). While this idea did not gain much traction in the decades after it was proposed, recent findings from our group and others have lent credence to this idea: We showed that the SC is a liquid crystalline material that remains dynamic after assembly (Rog et al., 2017). This evidence that proteins can move along the interface between paired chromosomes suggested a potential mechanism for communication between and patterning of recombination nodules (Zhang et al., 2018). A recent study proposed a mechanism based on diffusion and coarsening of an essential RN component along chromosomes (Morgan et al., 2021), similar to the model presented here.

Correlative and experimental evidence have indicated that the SC is essential for CO interference. Diverse evolutionary lineages, including some fungi and protists, have lost SCs, and their COs do not show interference. Budding yeast can make Class II COs in the absence of SC, but the interfering Class I COs are abolished (Sym, 1994); indeed, the “ZMM” proteins include Zip1, an essential component of the SC (Lynn et al., 2007). Perhaps most convincingly, *Arabidopsis* lacking the essential SC protein ZYP1 can still make crossovers via the “ZMM” pathway, but these do not show interference (Capilla-Pérez et al., 2021). In *C. elegans*, all crossovers are normally of the Class I type, and strictly depend on the SC (Colaiácovo et al., 2003; Kelly et al., 2000; Zalevsky et al., 1999). This evidence does not support a role for the SC in interference per se. However, reduced expression of SC components, or mutations that perturb the structure of the SC without eliminating its formation, can lead to increased numbers of LRNs and crossing-over, and a reduction in interference (Gordon et al., 2021; Köhler et al., 2020; Libuda et al., 2013).

Despite accumulating evidence that the SC plays a central role in crossover interference, this idea remains controversial. The primary counterevidence is that in *S. cerevisiae*, crossover-designated recombination intermediates arise during zygotene and act as SC initiation sites, yet the crossovers detected among meiotic products display interference (Börner et al., 2004).

Although our coarsening model relies on diffusion of pro-crossover proteins along the SC, which we demonstrate experimentally, the model also explains how interference can appear to precede SC assembly. Evidence from *Sordaria*, mice, *Drosophila*, and many plants, in addition to *C. elegans*, has indicated that recombination nodules undergo a progression from many early foci to a more limited number of “crossover-designated” foci (Wettstein et al., 1984), which our model attributes to coarsening. In *C. elegans* this transition is regulated by CHK-2 activity so that it normally occurs only after complete assembly of the SC and establishment of JMs along all chromosomes (Zhang et al., 2021). However, we postulate that in other organisms, including budding yeast, coarsening may occur concomitant with SC assembly. In these organisms, recruitment of pro-crossover factors to early recombination intermediates triggers initiation of synapsis at those sites. If competition for pro-crossover factors occurs as chromosomes synapse, recombination intermediates that arise later would not compete effectively and thus would rapidly be depleted of factors that protect them from noncrossover resolution. When two or more designated crossovers/SC initiation sites do arise in proximity to each other, one will win out once the SC spans the region between them.

By delaying coarsening and crossover designation until after the completion of synapsis, some systems may level the playing field to enable regions of the chromosomes that pair and synapse later – *e.g.*, sites far from telomeres or pairing centers, which can act as synapsis initiation sites, – to compete effectively for pro-crossover factors. Mechanisms that delay crossover designation until synapsis is completed also account for crossover assurance, since synapsis typically depends on the formation of JMs. In *C. elegans*, where synapsis does not depend on recombination intermediates, a “crossover assurance” checkpoint delays crossover designation until JMs are established on all chromosomes. Conversely, the absence of a pre-coarsening stage should favor CO designation at DSB sites close to telomeres or other early pairing sites, as is seen in many organisms. Differences in the timing of the early and late stages may also contribute to “heterochiasmy,” *i.e.*, differences in crossover distributions between male and female gametes in the same organism. Intriguingly, cytological evidence suggests that CO designation occurs before synapsis is completed in the nematode *Pristionchus pacificus*, yet the crossover distribution and interference in this organism are very similar to what is seen in C. elegans (Rillo-Bohn et al., 2021). Thus, it appears that similar crossover patterns can arise with or without a pre-coarsening stage.

We found that most conventional pathways that repress or activate CDKs are dispensable for crossover patterning. However, active CDK-2 becomes more concentrated in the late stage, leading to the growth of LRNs. During early pachytene, when many MSH-5 foci/ERNs are detected, most of the ZHP-3/4 remains associated with the SC rather than ERNs. This may limit the growth of individual nodules. In the late stage, ZHP-3/4 accumulates to a much greater extent in the nodules, which likely leads to an increase in positive feedback between CDK-2 activity and coarsening. A drop in CHK-2 activity may regulate this transition by influencing the partitioning of ZHP-3/4 between SC and RNs.

To understand how coarsening is regulated, we will need to investigate how CDK-2 activity is restricted in the early stage, and how it can be exchanged between RNs in the late stage. We speculate that ZHP-1/2, which are specifically required for coarsening in *C. elegans*, may be required to transport CDK-2 along the SC, while ZHP-3/4 are required for all CDK-2 activity. The confinement of these RING domain proteins to the SC, together with the fluid properties of the SC, can explain how exchange is limited to occur between RNs along the same pair of chromosomes.

### Have *C. elegans* recombination nodules been observed by electron microscopy?

A series of studies based on electron microscopy of *C. elegans* oocytes described nuclear structures associated with SCs. These fairly large (∼250 nm), granular bodies were first designated as “SC knobs” and later as “disjunction regulator regions (DRRs)” (Goldstein, 1986, 1982; Goldstein and Slaton, 1982). Six DRRs were detected in serial-section reconstructions of wild-type oocytes at pachytene, consistent with the crossover number and the six designated crossover sites detected by fluorescent microscopy. DRRs differ in appearance from RNs observed in *Drosophila*, plants, and even other nematodes; they show more internal heterogeneity and are somewhat larger than “typical” RNs. They were also reported to be absent from several crossover-proficient mutants such as *him-8* and *him-5,* casting doubt on their identification as recombination nodules. However, we now appreciate that the formation of LRNs is delayed until very late pachytene in these mutants due to meiotic checkpoints, which may account for the absence of DRRs in the few pachytene nuclei that were sectioned and analyzed in these studies. Correlatively light and electron microscopy should help to determine whether DRRs are indeed recombination nodules.

## Materials and methods

### Transgenic worm strains

All *C. elegans* strains were maintained under standard conditions on nematode growth medium (NGM) plates seeded with *E. coli* strain OP50 at 20°C. New alleles used in this study were generated by CRISPR/Cas9-mediated genome editing, as previously described (Zhang et al., 2018). To generate new *msh-5* alleles, repair templates were gel-purified and melted before injection (Ghanta and Mello, 2020).

To maintain strains and analyze the effects of mutations in *msh-5*, all of which caused strong meiotic defects, we crossed them to two different balancer chromosomes, *nT1* and *tmC9*. In some cases, the mutations underwent recombination with the balancer chromosomes, leading to the production of progeny that did not segregate the balancer but had a different phenotype from that observed in homozygotes when the mutations were first isolated. This was surprising, since these balancers are both very stable, and this may thus indicate a high frequency of aberrant recombination in heterozygous animals. To maintain these strains, we picked individual heterozygotes to plates and verified that their progeny showed Mendelian segregation of the balancer and *msh-5* mutant allele, which was also re-sequenced before our experiments. Brood counts and analysis of DAPI-staining bodies were performed using homozygous progeny of heterozygous parents, and heterozygotes carrying the *tmC9* balancer (Dejima et al., 2018), which did not affect segregation or progeny viability on its own, based on our analysis of GFP::MSH-5^WT^/*tmC9*.

### Brood analysis

To quantify brood size (number of embryos), male self-progeny, and embryo viability, L4 hermaphrodites were picked onto individual plates and transferred to new plates daily for 4 days. Embryos were counted after removing the parents, and re-counted later on the same day to increase the reliability of the counts. Viable progeny and males were counted when progeny reached L4 or adult stages.

### Protein depletion using auxin-induced depletion and/or RNA interference (RNAi)

Auxin-mediated protein depletion (AID) was performed as previously described (Zhang et al., 2018, 2015). Unless otherwise indicated, auxin (IAA, Alfa Aesar #A10556) was used at 1 mM. To combine AID depletion with RNAi, NGM agar was cooled to 55°C and supplemented with 0.5 mM or 4 mM auxin, 1 mM IPTG and 25 µg/ml carbenicillin just before pouring plates. RNAi plasmids for *cdk-2* knockdown were constructed by inserting a PCR product spanning exon 3 of *cdk-2* into plasmid vector pL4440, or exons 2-6 into plasmid vector T444T (Sturm et al., 2018) and transforming into *E. coli* strain HT115. Cultures containing the RNAi plasmid or the corresponding empty vector were cultured overnight at 37°C in the presence of 25 µg/ml carbenicillin and 1mM IPTG to induce dsRNA synthesis. The culture was then concentrated 50-fold, supplemented with auxin at the same concentration as on the plates, or ethanol (solvent for IAA) and spread on plates. Hermaphrodites at the L4 stage were picked, washed twice with S-basal and transferred to RNAi ±auxin plates. Two hours later worms were transferred to new RNAi ±auxin plates and incubated at 20°C. Although *cdk-2(RNAi)* did not dramatically decrease CDK-2 on its own, it enhanced the penetrance of AID-mediated depletion.

### Construction of an analog-sensitive CDK-2 allele and inhibitor analysis

Sequence alignments identified phenylalanine 91 in CDK-2 as the “gatekeeper” residue (Lopez 2014). Using CRISPR/Cas9 we mutated to either glycine or alanine. Both strains were tested for effects on viability and/or germline proliferation when grown on plates containing one of five different bulky ATP analogs: 3-IB-PP1, 3-MB-PP1, 1-NA-PP1, 3-BrB-PP1, and 1-NM-PP1. These compounds were dissolved in 100% ethanol and added to NGM immediately before pouring plates. OP50 *E. coli* grown overnight in LB was concentrated by centrifugation and resuspended in minimal media. 40 µl of concentrated OP50 was pipetted onto each plate and allowed to dry overnight at 4**°**C. No effects were observed in wild-type animals grown on plates containing these inhibitors at concentrations up to 40 µg/ml. Hermaphrodites homozygous for the F91G mutation showed sensitivity to several inhibitors, while the F91A allele was not analog-sensitive. A concentration of 8 μg/ml 3-IP-PP1 was sufficient to prevent reproduction in the *cdk-2^F91G^* strain (hereafter *cdk-2^as^*).

### Fluorescent labeling of replicating DNA

Adult worms were picked from plates into M9 buffer containing OP50 bacteria and 1mM 5-ethynyl-2′-deoxyuridine (EdU). Following a 15-minute incubation, animals were washed once with egg buffer and then immediately dissected for staining. Fixation was carried out as for immunofluorescence except that 3% formaldehyde was used and fixation was performed for 30 minutes with rotation. Fixative was replaced with prechilled methanol for 15 min. Samples were washed two times with PBST, blocked for 1 hour and incubated with primary antibodies overnight at 4°C. After 3 washes, secondary antibodies were applied for 1 hour at room temperature. Labeling with Alexa Fluor 488 azide was performed in a 250-µl reaction according to the manufacturer’s recommended procedure (Thermo Fisher Catalog #C10637) for 30 min in the dark at room temperature with rotation. Samples were washed 3 times before mounted in Prolong Diamond mounting medium with DAPI (Life Technologies #P36962).

### Yeast two-hybrid analysis

Interactions between CDK-2 and COSA-1 were assayed using the Matchmaker Gold Yeast Two-Hybrid System (Clontech PT4084-1). CDK-2 and COSA-1 cDNAs were amplified from a *C. elegans* cDNA library. Each coding sequence was fused to the GAL4 DNA binding domain and GAL4 activation domain by insertion into pGBKT7 and pGADT7, respectively, using Gibson assembly (New England Biolabs #M5510A). Plasmids were transformed into the Y2HGold yeast strain using standard lithium acetate-mediated transformation. Positive and negative controls provided with the kit were included in all assays.

### Immunoprecipitation

Animals were allowed to starve on standard 60-mm worm plates to obtain large populations of synchronized L1 larvae. 30 plates for each genotype were washed to collect the animals, which were concentrated and then transferred to 500 mL liquid cultures. They were grown with aeration at 200 rpm at 20°C for 3 days, when they reached adulthood. Worms were then collected and frozen in liquid nitrogen. The frozen pellets were processed using a mixer mill (Retsch MM40) to disrupt the cuticle, thawed by addition of cold lysis buffer (25 mM HEPES pH7.4, 100 mM NaCl, 1 mM MgCl_2_, 1 mM EGTA, 0.1% Triton X-100, 1 mM DTT, phosSTOP (Sigma #4906837001) and cOmplete protease inhibitors (Sigma #4693159001), and sonicated using a Branson Digital Sonifier while on ice. The resulting lysate was centrifuged at 20,000 RCF for 20 min at 4°C. Supernatant was collected and incubated with anti-FLAG M2 magnetic beads (Sigma #M8823) for 3 hours at 4°C. Beads were then washed 4 times with lysis buffer, and proteins were eluted using a 3xFLAG peptide (Sigma #F4799). Samples were then heated to 100°C for 10 minutes in SDS sample buffer and analyzed by western blotting.

### Western blotting

Adult worms were picked into S-basal buffer and lysed by adding an equal volume of 2X SDS sample buffer and heating to 100°C for 30 minutes with intermittent vortexing. Whole worm lysates were then separated on 4-12% polyacrylamide gradient gels (GenScript #M00652 and #M00654), transferred to nitrocellulose membranes, and blotted with indicated antibodies: Mouse anti-FLAG (1:1,000, Millipore Sigma, #F1804), Mouse anti-V5 (1:1,000, Thermo Fisher, #R960-25) and Mouse anti-α-tubulin (1:5,000, Millipore Sigma, #05–829). HRP-conjugated anti-mouse secondary antibodies (Jackson Immunoresearch #115-035-068) and SuperSignal™ West Femto Maximum Sensitivity Substrate (Thermo Fisher #34095) were used for detection.

### Immunofluorescence

Most immunofluorescence experiments were performed on slides as previously described (Zhang et al., 2018). Images shown in Figures 1, 3, and S1 were obtained from dissected animals stained in1.5-ml tubes. Briefly, young adult hermaphrodites (20-24 h post-L4) were dissected in Egg Buffer containing 15 mM sodium azide and 0.1% Tween 20, followed by fixation with 1% formaldehyde in the same buffer on a coverslip for 2 min. The worms were then transferred to PBS containing 0.1% Tween 20 (PBST) for 5 min. PBST was aspirated and replaced with methanol pre-chilled to −30°C for 5 min. Samples were then washed 3 times for 5 min each in PBST and blocked with 1X Blocking Reagent (Roche) in PBST. Primary antibody incubations were performed overnight at 4°C. After 3 washes with PBST, secondary antibody (including fluorophore conjugated nanobodies) incubations and DAPI staining were conducted sequentially at room temperature. Samples were then mounted with SlowFade Glass mounting medium (Invitrogen) using High Precision #1.5 coverslips (Zeiss/Marienfeld).

Primary antibodies were purchased from commercial sources or have been previously described. They were used at the following dilutions: Mouse anti-FLAG (1:500, Sigma, #F1804), Mouse anti-HA (1:400, Thermo Fisher, #26183), Mouse anti-V5 (1:500, Thermo Fisher, #R960-25), Cy3 conjugated FluoTag®-X2 anti-ALFA, (1:500 NanoTag # N1502-SC3), Mouse anti-GFP (1:500, Millipore Sigma, #11814460001). Rabbit anti-H3pSer10 (1:500, EMD Millipore, #382159), Rabbit anti-SYP-2 [1:500, (Colaiácovo et al., 2003)], Rabbit anti-RAD-51 (1:5,000, Novus Biologicals, #29480002), Goat anti-SYP-1 [1:300, (Harper et al., 2011)], Chicken anti-HTP-3 [1:500, (MacQueen et al., 2005)], Rat anti-HIM-8 [1:500, (Phillips et al., 2005, p. 8)], Secondary antibodies labeled with Alexa 488, Cy3, or Cy5 were purchased from Jackson ImmunoResearch (WestGrove, PA) and used at 1:500. Images were acquired using a DeltaVision Elite microscope (GE) equipped with a 100x, 1.4 N.A. or 100x 1.45 N.A. oil-immersion objective. Z-stacks of optical sections spanning 8-12-µm were collected at 0.2 µm z-spacing. Iterative 3D deconvolution and image projections were performed using the SoftWoRx package; colorization was done using Adobe Photoshop 2021.

### Quantification of univalent and bivalents at diakinesis

To count DAPI-staining bodies in *msh-5* mutants, hermaphrodites were aged 2-3 days from the L4 stage. Animals were picked into a small drop of water on a HistoBond slide. Most of the water was removed by touching the drop with a piece of Whatman filter paper. Worms were then fixed and dehydrated by dropping a large drop of 95% ethanol from a Pasteur pipet directly onto them, followed by air-drying. Ethanol addition and air drying was repeated 1-2 additional times. The air-dried sample was then mounted by addition of glycerol-based mounting medium containing 0.5 µg/ml DAPI and imaged using a DeltaVision Elite microscope using a 100X NA 1.4 oil immersion objective.

### Imaging of MSH-5 nonphosphorylatable and phosphomimetic mutants

Mutant alleles were generated in a strain expressing GFP::MSH-5. To confirm the expression of the mutant proteins, adult hermaphrodites (24 h post-L4) homozygous for each allele were picked into a solution of 60% PBS, 20% glycerol, and 20% PEG 20,000 (v/v), covered with a coverslip and sealed with VALAP (1:1:1 petroleum jelly:lanolin:paraffin). Worms were imaged immediately after mounting using a Marianas spinning-disc confocal microscope (3i) equipped with a 100× 1.46 NA oil immersion objective. Images were acquired under the same conditions for each mutant (100% laser power, 250 ms exposure time) and scaled identically to facilitate comparison.

### Fluorescence recovery after photobleaching (FRAP)

All FRAP imaging was performed on age-matched young adults, 24-hour post-L4. Hermaphrodites expressing a fluorescently tagged protein of interest were picked into a 5-µl drop of M9 buffer containing 0.4% tricaine and 0.04% tetramisole on top of a thin agarose pad. These were covered with a #1.5 coverslip and sealed with VALAP. Regions containing late pachytene nuclei that were near the surface of the worm and near the coverslip were selected for analysis.

Photobleaching and imaging were performed on a Marianas spinning-disc confocal microscope from 3i (Intelligent Imaging Innovations, Inc.) equipped with a 100X 1.46 NA oil immersion objective and vector photomanipulation module. Photobleached areas were pre-selected using the rectangle selection tool in Slidebook 6. Photobleaching was performed with the 488nm laser at 100% laser power. Z-stacks centered around the plane of focus during photobleaching were collected at 0.25 µm spacing spanning a total thickness of 11 µm. Images were acquired at 50% laser power using 100 ms exposures. For each photobleaching experiment, a time series was collected at 1-minute intervals, including one reference time point collected before photobleaching and twenty-two subsequent images, for a total acquisition time of twenty-four minutes. Additive projections of the 3D data stacks for each time point were generated using ImageJ and stored as TIF images. Background subtraction was performed using the “Subtract Background” function with a rolling ball size set to 5 pixels. Photobleached foci were identified as ROIs (regions of interest) using the “Auto Threshold” function of ImageJ set to “Default,” then increasing the threshold until no diagonally associated pixels were included in each ROI. If any significant movement of foci occurred, ROIs were manually realigned. Average pixel intensities for each ROI were normalized by the corresponding prebleach value to measure the relative fluorescence.

### FCS immobilization, imaging, and analysis

All FCS experiments were performed on young adult hermaphrodites 24h after the L4 stage. Immobilization and sample preparation were performed as previously described using polystyrene beads and serotonin (Rog et al., 2017), with the addition of 0.2% tricaine and 0.02% tetramisole in the mounting medium. Regions of the germline that were close to the coverslip were used for imaging and analysis.

Most of the line-scan data for ACF and pCF analysis was performed on a Zeiss LSM 710 with a 63X 1.4 NA oil immersion objective. Images for ZHP-1-4 labeled with split-GFP, and controls with ZHP-3::GFP, were acquired using a Zeiss LSM 880 with a 63X 1.4 NA oil immersion objective. Microscopy settings for the LSM 710 were as follows. The pinhole was set to acquire voxels spanning 1.1 µm in z. Voxel dimensions in the XY plane were set using the “optimal setting” function on a line-scan selection with zoom of 125X (1.08 µm line length). This selected eight voxels each 0.135 µm in length. The duration of each scan was 473 µs, and 1000 successive scans were collected at maximum speed. For scans acquired using the LSM 880, the pinhole was set to acquire voxels 0.8 µm in z, a zoom of 100X (1.35 µm line length) was used, and optimal setting selected sixteen voxels, each 0.0844 µm in length. Each line acquisition took 292 µs; 40,000 lines were collected per scan for a total imaging time of 11.68 s. The “smart setup” filter sets for EGFP were used, with 2% laser power, for both sets of data.

The autocorrelation function (ACF) of each voxel was calculated in Matlab, using previously described equations (Wachsmuth et al., 2000). The ACF for individual voxels as well as average ACF for entire line-scans (LSM 710) or half line-scans (LSM 880) were analyzed, which increases the total time (Qian and Elson, 1991). A fluorescein solution was used to determine the value of *w*_0_ (Digman and Gratton, 2009), which was found to have an average value of 0.29 µm (LSM 710) and 0.10 µm (LSM 880). The ACF data were best fit to a two-dimensional diffusion model (Vukojević et al., 2005), which was applied to estimate diffusion coefficients using a least-squares fit algorithm to the average ACF of a scan/half scan (Figure 5), or for every voxel in every scan (Figure S9). SimFCS software was used to calculate the pCF (Digman and Gratton, 2009) for voxel pairs at a distance of 0.135 µm (LSM 710) or 0.165 µm (LSM 880). The correlation time that gives the maximum pCF indicates the average time it takes for a fluorescent particle to move that given distance. We extracted these times from the pCFs and report them as the “movement index” for each scan/half-scan (Figure 5) or for each voxel pair (Figure S9).

The autocorrelation of fluorescence intensity within a specified volume – i.e., a voxel – decays over time due to the movement of fluorophores into and out of that volume. Diffusion rates can thus be estimated from ACFs. Because this relationship depends on the imaging parameters, estimates of diffusion rates based on ACF analysis are calibrated using a fluorophore with a known diffusion rate; we used a solution of fluorescein. This enabled us to fit a diffusion model and to estimate diffusion coefficients for fluorescent proteins labeled with GFP or split-GFP, based on the average of ACFs calculated for each scan (Figure 5D). In parallel, we estimated diffusion coefficients based on ACFs for all voxels in all scans for each protein (Figure S9A), which yielded very similar values.

### Multiple sequence alignment

Predicted MSH-5 proteins from *Caenorhabditis* species were obtained from (http://download.caenorhabditis.org/v1/json/CGP_orthology.html) and edited by removing sequences that were incomplete or contained large insertions relative to others. The multiple sequence alignment shown in Figure S8 was calculated using MAFFT (Katoh et al., 2019) and displayed using Jalview (Waterhouse et al., 2009).

### Quantification and statistical analysis

Quantification methods and statistical parameters are described in the legend of each figure, including *n* values, error calculations (SD or SEM), statistical tests, and *p*-values. *p*<0.05 was regarded as significant.

## Acknowledgements

Some strains used in this work were provided by the CGC, which is funded by NIH Office of Research Infrastructure Programs (P40 OD010440). We thank members of the Dernburg lab for helpful discussions during the course of this work.

## Author Contributions

Conceptualization, L.Z. W.S. D.Z. and A.F.D.; Methodology, L.Z. W.S. D.Z. and A.F.D.; Investigation, L.Z. W.S. and A.F.D.; Writing – Original Draft, L.Z. W.S. D.Z. and A.F.D.; Writing – Review & Editing, L.Z. W.S. D.Z. and A.F.D.; Funding Acquisition, D.Z. and A.F.D.; Resources, L.Z. W.S. D.Z. and A.F.D.; Supervision, D.Z. and A.F.D

---

**Figure S1.**
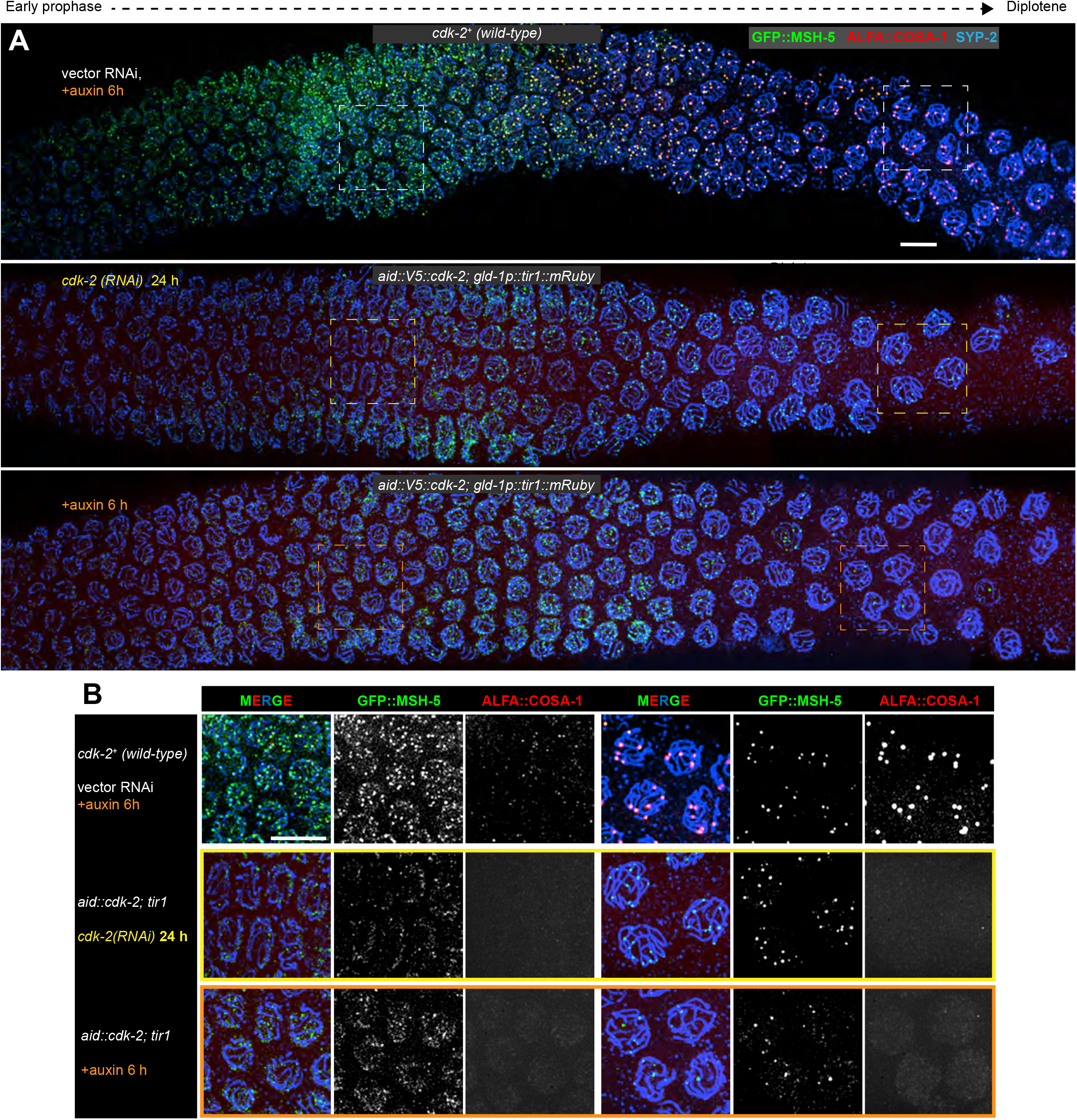
Incomplete depletion of CDK-2 by RNAi or AID alone, Related to Figure 1. (A) Immunolocalization of GFP::MSH-5 (green), ALFA::COSA-1 (red) and SYP-2 (blue) in a hermaphrodite germline. Upper panel: recombination intermediates are not affected by auxin treatment of a control strain expressing CDK-2 without a degron. *cdk-2(RNAi)* treatment for 24 hours or 4mM auxin exposure for 6 hours greatly reduce but do not abolish recombination intermediates. ALFA::COSA-1 localization was more sensitive to CDK-2 depletion than GFP::MSH-5. These samples were placed on RNAi or auxin plates and stained in parallel with those shown in Figure 1E-F. Scale bars, 5 µm. (B) Enlargements of the indicated regions from (A). Scale bars, 5 µm.

**Figure S2.**
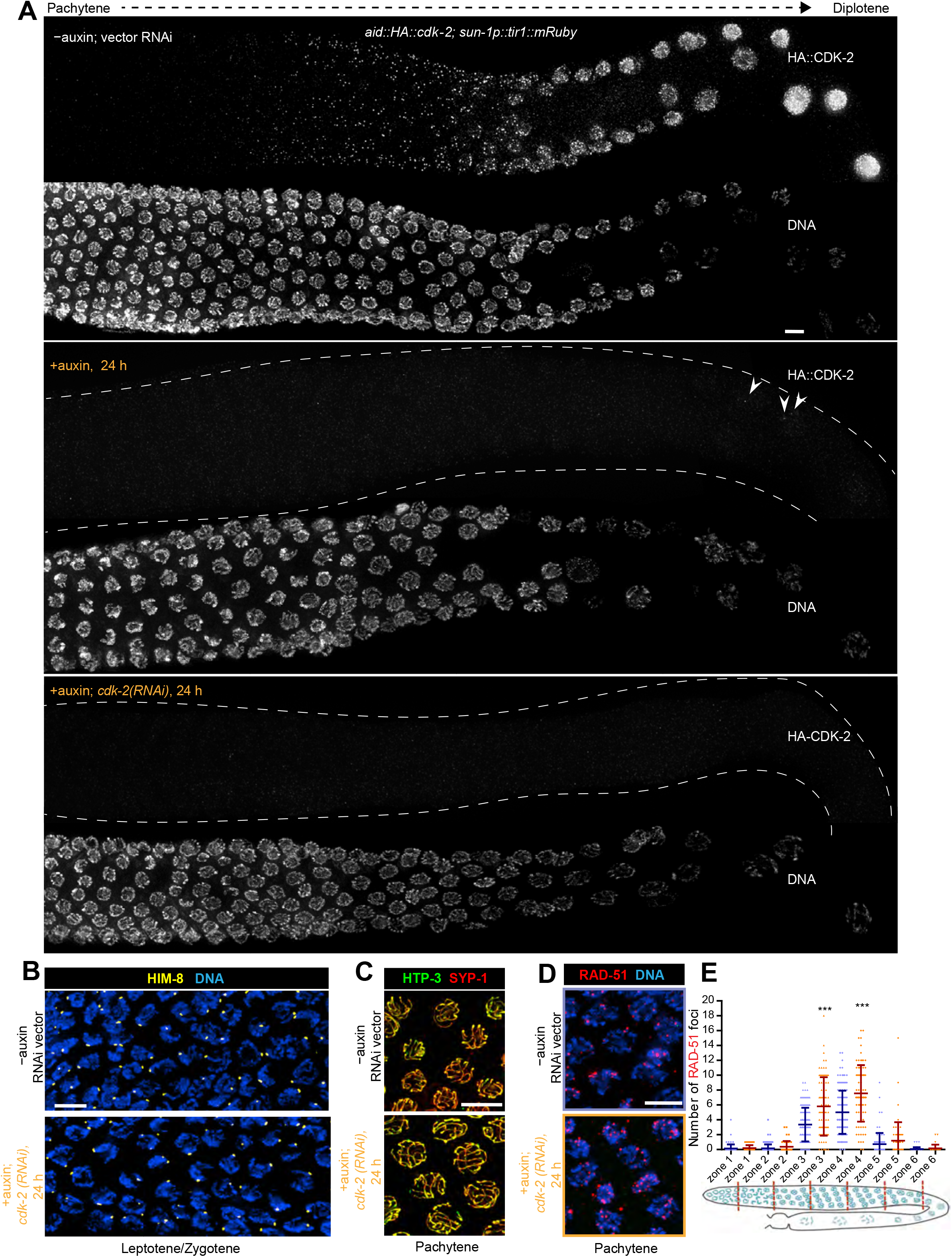
CDK-2 is dispensable for homolog pairing, synapsis and DSB induction, Related to Figure 2. (A) Late prophase in *aid::HA::cdk-2; sun-1p::tir1* hermaphrodites stained with anti-HA antibodies to confirm effective depletion of CDK-2 by RNAi + auxin treatment. Scale bars, 5 µm. (B) Early prophase nuclei in *aid::HA::cdk-2; sun-1p::tir1::mRuby* gonads stained for HIM-8 (yellow), which marks the pairing center region of the X chromosome (Phillips et al., 2005), and DNA (blue). Robust pairing of HIM-8 foci is observed following depletion of CDK-2. Scale bars, 5 µm. (C) Nuclei at mid-prophase stained for HTP-3 (green) and SYP-1 (red). Colocalization of these proteins reveals formation of the SC along all axes in controls and hermaphrodites depleted of CDK-2, as described in (A). Scale bars, 5 µm. (D) Nuclei at mid-prophase stained for RAD-51 (red) and DNA (blue), showing abundant DSB repair intermediates marked by RAD-51 following depletion of CDK-2 as described in (A). Scale bars, 5 µm. (E) Dynamics of RAD-51 foci. Hermaphrodite gonads were divided into six zones of equal length spanning the premeiotic through late pachytene region, as shown in the diagram below. The number of RAD-51 foci in each nucleus is shown for each zone, with mean and SD values indicated by bars. 3 gonads were scored for each condition. Nuclei counted for each zone were as follows: for controls – 74, 113, 113, 127, 86, 47; for CDK-2 depletion – 47, 46, 80, 82, 58, 33. More nuclei were present in the controls due to effects of CDK-2 depletion on germline proliferation. ***p<0.0001, Mann-Whitney test.

**Figure S3.**
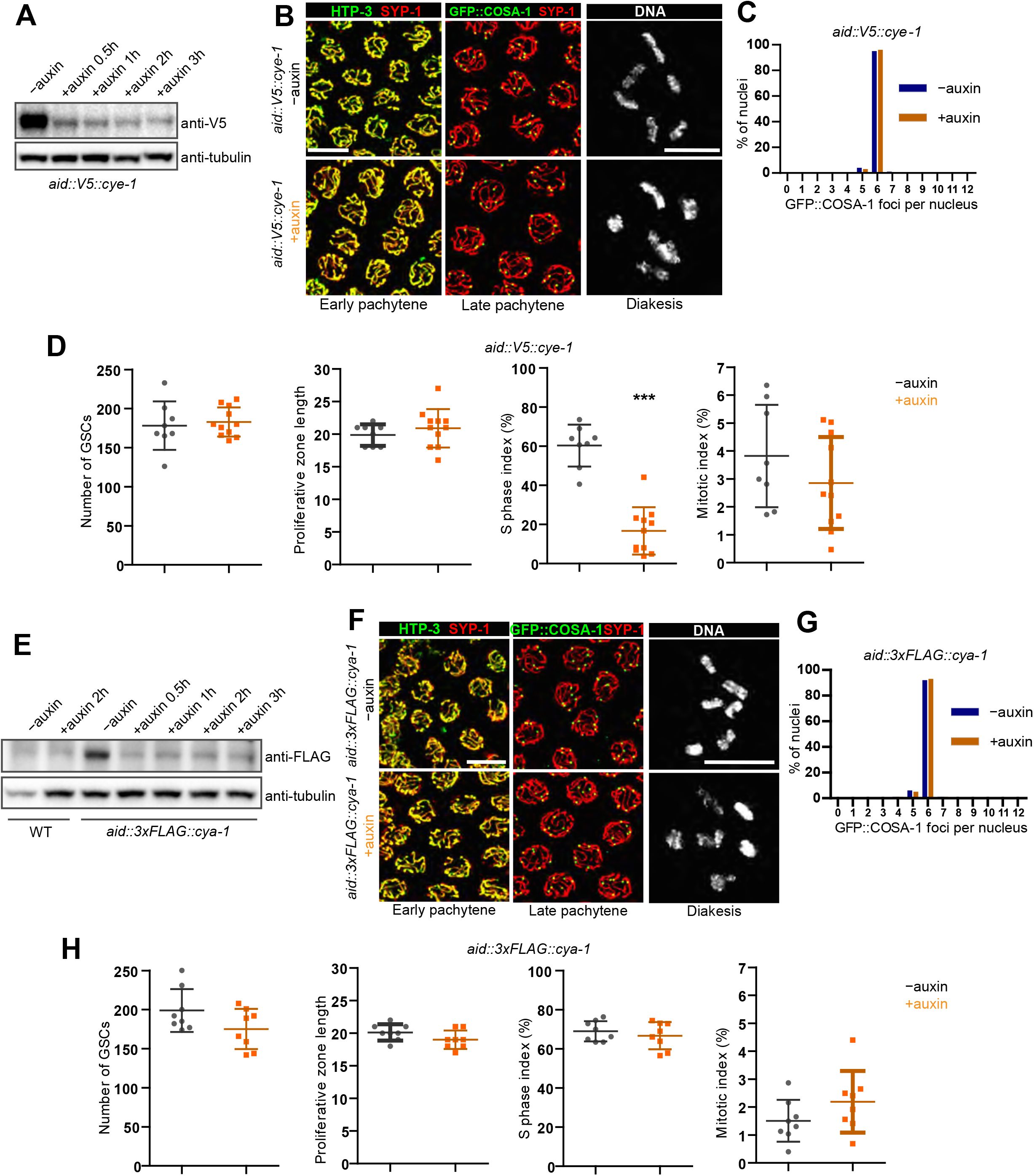
Cyclin E and cyclin A are dispensable for crossing-over, Related to Figure 2. (A) Western blot of lysates from worms expressing epitope-tagged cyclin E (AID::V5::CYE-1), detected with anti-V5 antibodies. Quantification of bands indicated > 95% reduction after 3h of auxin treatment. Tubulin was blotted as a loading control. (B) Depletion of CYE-1 does not affect synapsis (left panels), crossover designation (center panels), or bivalent formation. Hermaphrodites were treated with or without auxin for 24 hr. Scale bars, 5 µm. (C) Quantification of GFP::COSA-1 foci in late pachytene nuclei, displayed as a histogram. Seven gonads were scored for each condition (no auxin or auxin treatment for 24h). *p*=0.9914 by Chi-square test for trend. n = 276 (-auxin control) and 241 (+auxin) nuclei. (D) Depletion of cyclin E (CYE-1) affects germline From left to right: the number of germline stem cells (GSCs) per gonad, the length of proliferative zone, the percentage of cells in S phase and the percentage of mitotic cells. Data were derived from eight and eleven gonads for -auxin controls and 24-h depletion, respectively, and presented as mean ±SD. p=0.6934 and 0.3848, p<0.0001 and *p*=0.2439, respectively, two-sided Student *t*-test. S phase index was dramatically reduced in the pre-meiotic region upon CYE depletion. (E) Western blots of whole worm lysates from wild-type controls and animals expressing degron- and epitope-tagged cyclin A (AID::3xFLAG::CYA-1) showing effective depletion by auxin treatment. Tubulin was blotted as a loading control. (F) Images of representative pachytene nuclei and oocyte nuclei stained for HTP-3 or GFP-COSA-1 (green) and SYP-1 (red), or DNA (grey), indicating normal synapsis, crossover designation and chiasmata formation in the absence of CYA-1. Worms were treated with or without auxin for 24 hr. Scale bars, 5 µm. (G) Quantification of GFP::COSA-1 foci in late pachytene nuclei following depletion of CYA-1. Seven gonads were scored for each condition; 6 rows of nuclei were scored in each gonad.. P=0.8237 by Chi-square test for trend. n = 268 (-auxin control) and 286 (+auxin) nuclei. (H) Effects of CYA-1 depletion on GSCs. Bars represent mean and SD. *p*=0.0958, 0.1136, 0.4695 and 0.1691, respectively, by two-sided Student *t*-test.

**Figure S4.**
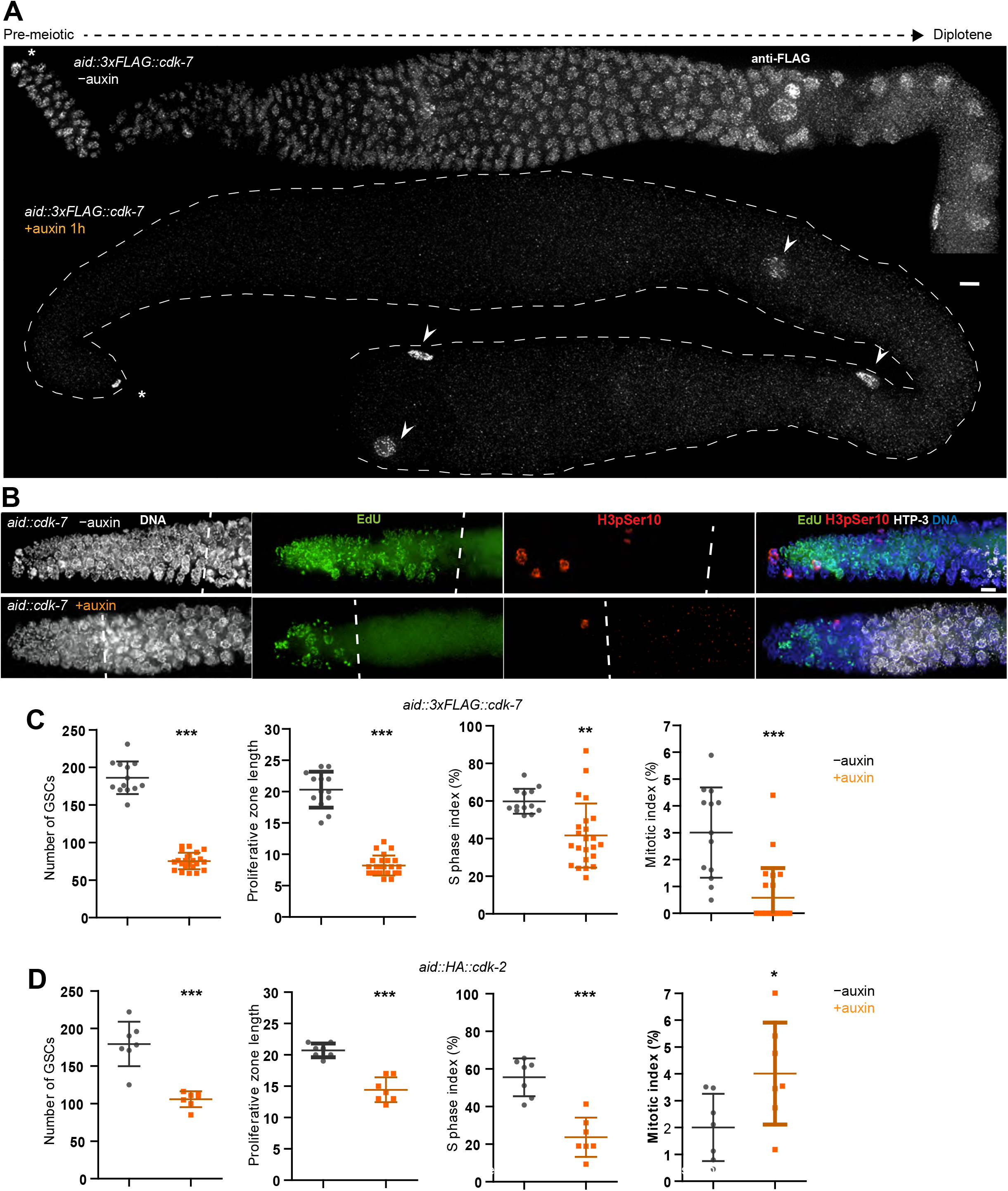
Depletion of CDK-7 causes cell cycle arrest in GSCs, Related to Figure 2. (A) Hermaphrodite gonads stained with anti-FLAG antibodies reveal efficient depletion of AID::3xFLAG::CDK-7 throughout the germline following treatment with 1 mM auxin for 1h. Asterisks indicate the distal tip cell and arrowheads point to somatic gonadal sheath cells, which do not express TIR1 in this strain, and CDK-7 is therefore not degraded. Scale bars, 5 µm. (B) GSCs are greatly reduced following CDK-7 depletion for 24 h. Dashed lines indicate boundaries between GSCs and meiotic entry, which is detected by localization of HTP-3 to chromosome axes. Scale bars, 5 µm. (C) Quantification of GSC number and cell cycle distribution following depletion of CDK-7 for 24h. Each data point represents one gonad; data were derived from 13 and 23 gonads, respectively. Data are presented as mean ± SD. \*\**p*=0.0008 and \*\*\**p*<0.0001, respectively, by two-sided Student’s *t*-test. (D) Data shown in Figure 2, panel F for CDK-2 depletion are repeated here to facilitate comparison with CDK-7 depletion. Surprisingly, depletion of CDK-7 had a stronger effect on GSC proliferation. This may indicate some redundancy between CDK-2 and CDK-1 for mitosis in GSCs.

**Figure S5.**
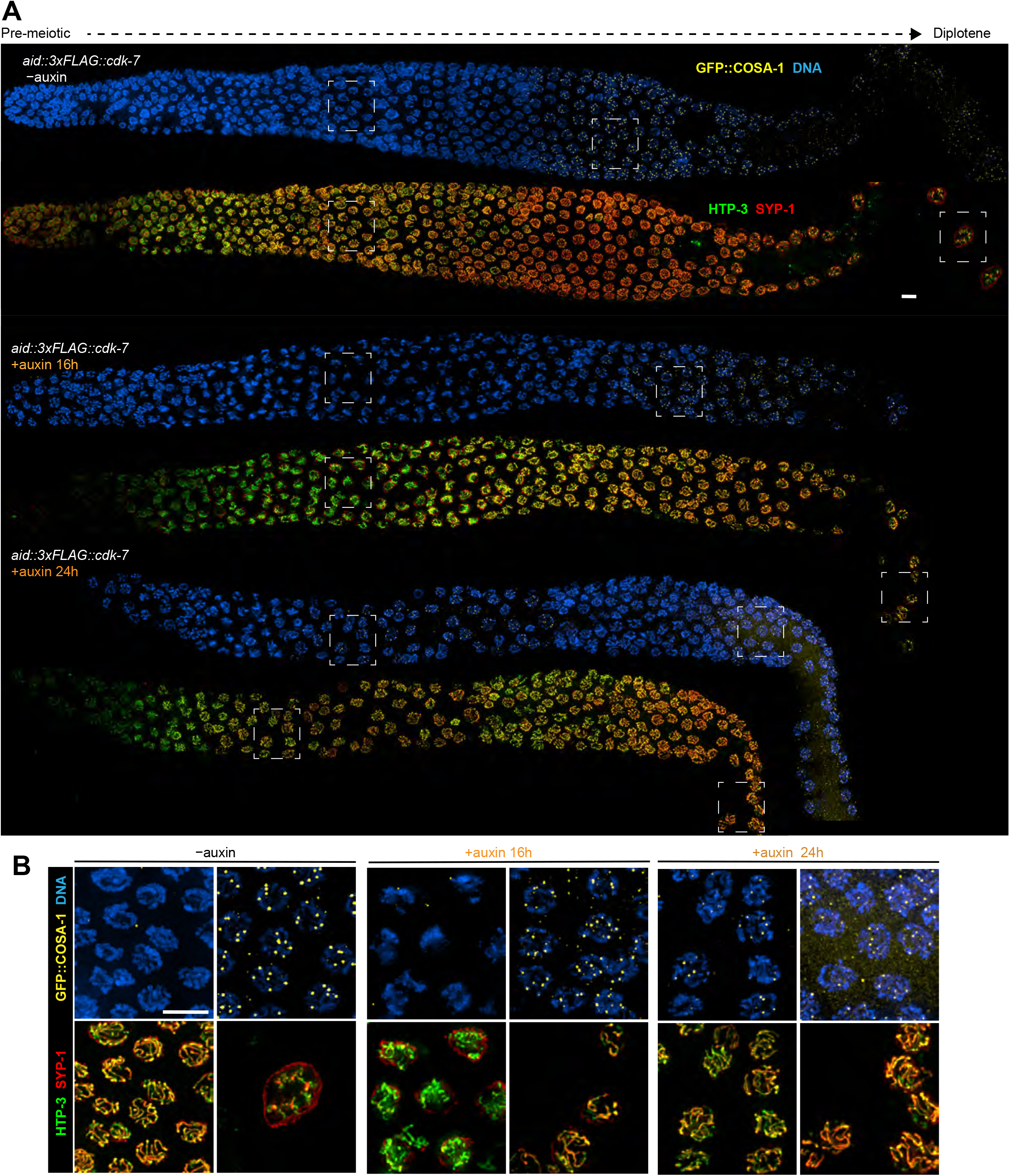
Depletion of CDK-7 perturbs meiotic progression. (A) Germlines from *aid::3xFLAG::cdk-7; sun-1p::tir1* hermaphrodites stained for HTP-3 (green), SYP-1 (red), GFP-COSA-1 (yellow) and DNA (blue). Worms were treated with or without 1mM auxin for 16 h or 24 h before analysis. As highlighted in the enlarged images, depletion of CDK-7 depletion results in delayed synapsis and failure to form a single row of cellularized oocytes following the bend in the gonad. Nevertheless, crossover designation occurs robustly. Scale bars, 5 µm. (B) Higher magnification of nuclei in the boxed areas in (A). Scale bars, 5 µm.

**Figure S6.**
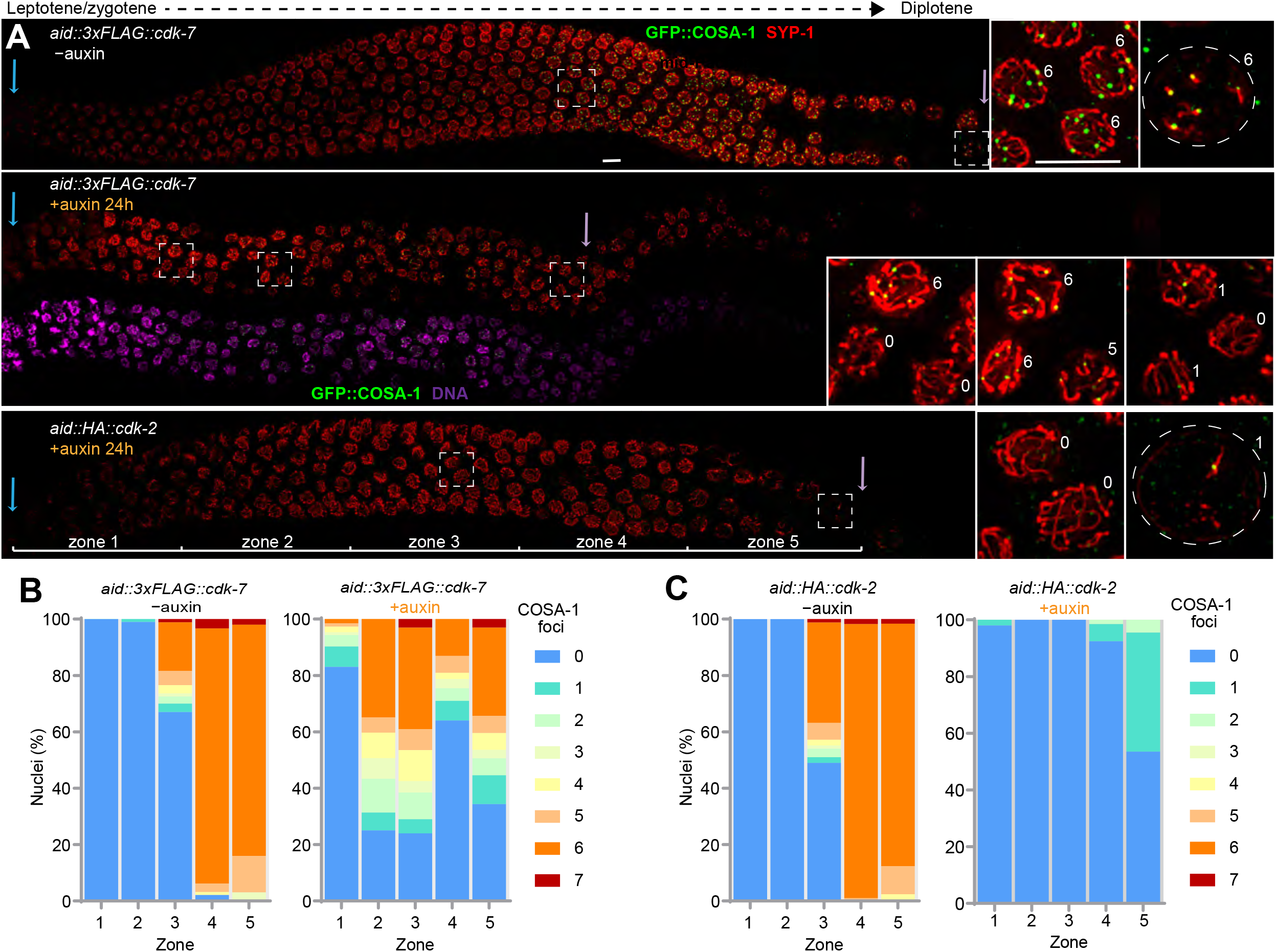
CO designation is perturbed but not abrogated by depletion of CDK-7, Related to Figure 2. (A) Germlines stained for GFP-COSA-1 (green), SYP-1 (red), and DNA (magenta, middle panel), reveal that crossover designation (appearance of COSA-1 foci) is perturbed in CDK-7, but completely abrogated by depletion of CDK-2. Hermaphrodites homozygous for degron-tagged CDK-7 or CDK-2 were treated with or without 1mM auxin for 24 h. Although CDK-7 depletion leads to more severe phenotypes in the pre-meiotic region than CDK-2 depletion (Figures 2 and S4), depletion of CDK-2 has a much more penetrant effect on crossover designation. Loss of CDK-7 may indirectly affect crossover formation – e.g., by delaying synapsis. Boxed areas indicate nuclei shown at higher magnification to the right. Blue arrows indicate meiotic entry. Purple arrows indicate the end of pachytene region. Scale bars, 5 µm. (B-C) Quantification of GFP::COSA-1 foci following depletion of CDK-7 or CDK-2. Hermaphrodite gonads were divided into five zones of equal length, spanning the meiotic entry through late pachytene region as shown in the (A) at the bottom. Data for CDK-7 depletion were derived from 5 control and 7 auxin-treated gonads; data for CDK-2 were from 4 control and 4 auxin-treated samples. Distributions of COSA-1 foci within each zone represent 40-300 nuclei.

**Figure S7.**
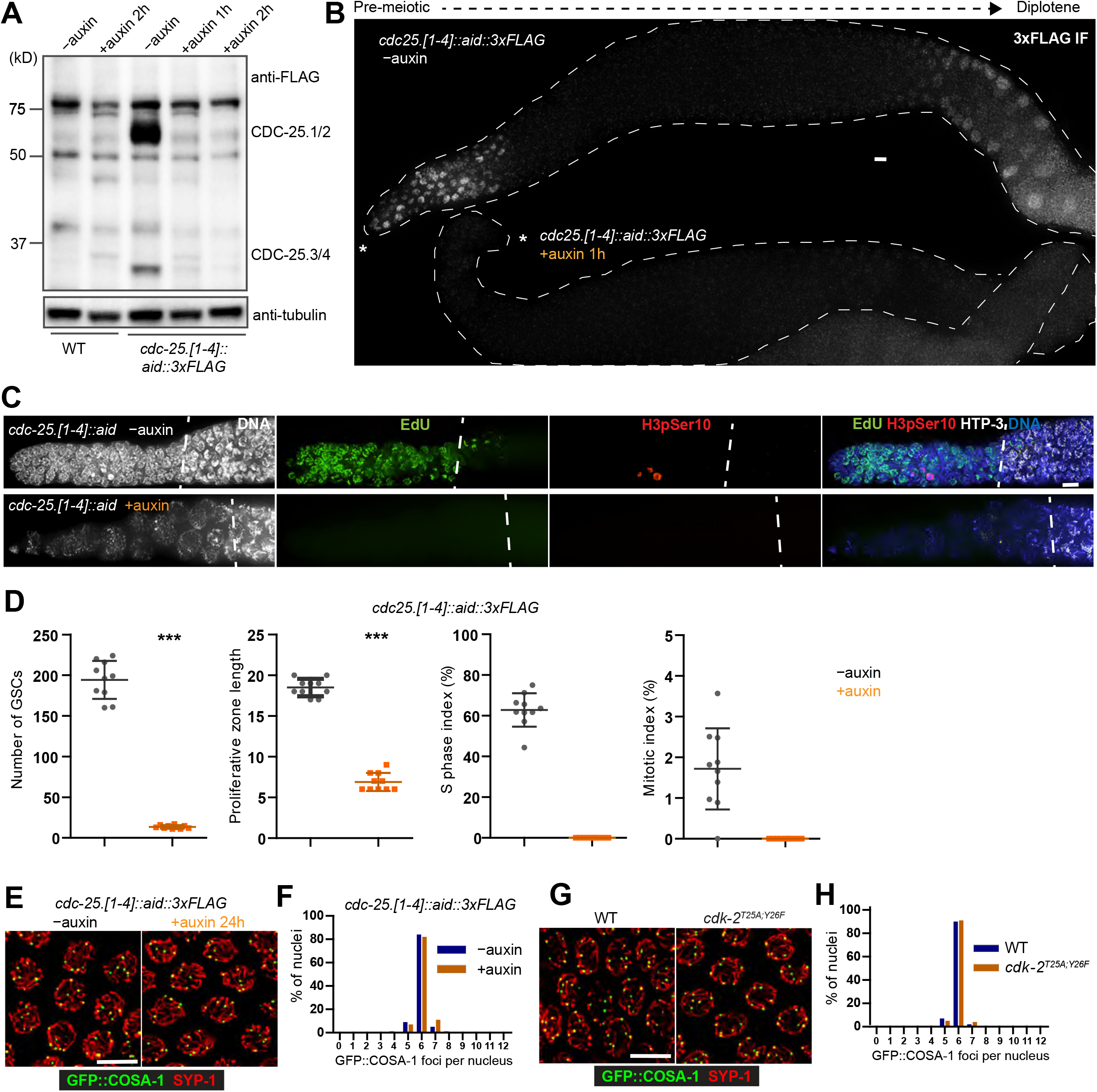
CDC-25 and N-terminal modification of CDK-2 are dispensable for crossing over, Related to Figure 2. (A) Western blots of total protein from wild-type hermaphrodites and a strain in which all four endogenous *cdc-25* genes (*cdc-25.1, cdc-25.2, cdc-25.3* and *cdc-25.4*) were tagged with a degron and 3xFLAG epitope. Proteins were detected with anti-FLAG antibodies. CDC-25.1 is the most abundant of the four proteins in young adults, consistent with quantitative proteomic studies (Grün et al., 2014). Robust depletion was detected within 1 hr of auxin exposure. Wild-type worms were treated and blotted in parallel as controls. Tubulin was blotted as a loading control. (B) Hermaphrodite gonads from the quadruple CDC-25 degron/FLAG-tagged strain, stained with anti-FLAG antibodies. Following 1 mM auxin treatment for 1 hr, CDC-25 proteins are not detected by immunofluorescence in germline nuclei. Scale bars, 5 µm. (C) GSC proliferation is arrested by depletion of CDC-25. Dashed lines indicate the boundary between pre-meiotic GSCs and meiotic prophase nuclei, which are identified by the presence of HTP-3 along chromosome axes. Worms were treated with or without 1 mM auxin for 24 h, followed by 15 min EdU incorporation and immunostaining. H3pSer10-positive nuclei are in mitosis. Scale bars, 5 µm. (D) Quantification of proliferative defects in GSCs following CDC-25 depletion. Animals were treated with or without 1 mM auxin for 24 h, followed by 15 min EdU corporation fixation and staining. Each data point represents one gonad; 10 were analyzed for each condition. Bars indicate mean and SD. ***p<0.0001, two-sided Student *t*-test. No cells in S-phase or mitosis were detected following depletion of CDC-25. (E) Images of representative late pachytene nuclei stained for GFP-COSA-1 (green) and SYP-1 (red), indicating normal crossover designation in the absence of CDC-25. Worms with four CDC-25 homologs (*cdc-25.1, cdc-25.2, cdc-25.3* and *cdc-25.4*) tagged with AID together with epitope 3xFLAG were treated in the presence or in the absence of auxin for 24 h before analysis. Scale bars, 5 µm. (F) Graphs indicating the distribution of GFP-COSA-1 foci in late pachytene nuclei in worms treated as in (E). P= 0.2059 by Chi-square test for trend. n is the number of nuclei scored for each condition. n= 253 and 320, respectively. Six and ten gonads were scored respectively. (G) Images of representative late pachytene nuclei stained for GFP-COSA-1 (green) and SYP-1 (red), indicating normal crossover designation in *cdk-2^T25A Y26F^* mutants. Scale bars, 5 µm. (H) Graphs indicating the distribution of GFP-COSA-1 foci in late pachytene nuclei in worms treated as in (G). P=0.3397 by Chi-square test for trend. n is the number of nuclei scored for each condition. n= 548 and 420, respectively. Eleven gonads were scored for each genotype.

**Figure S8.**
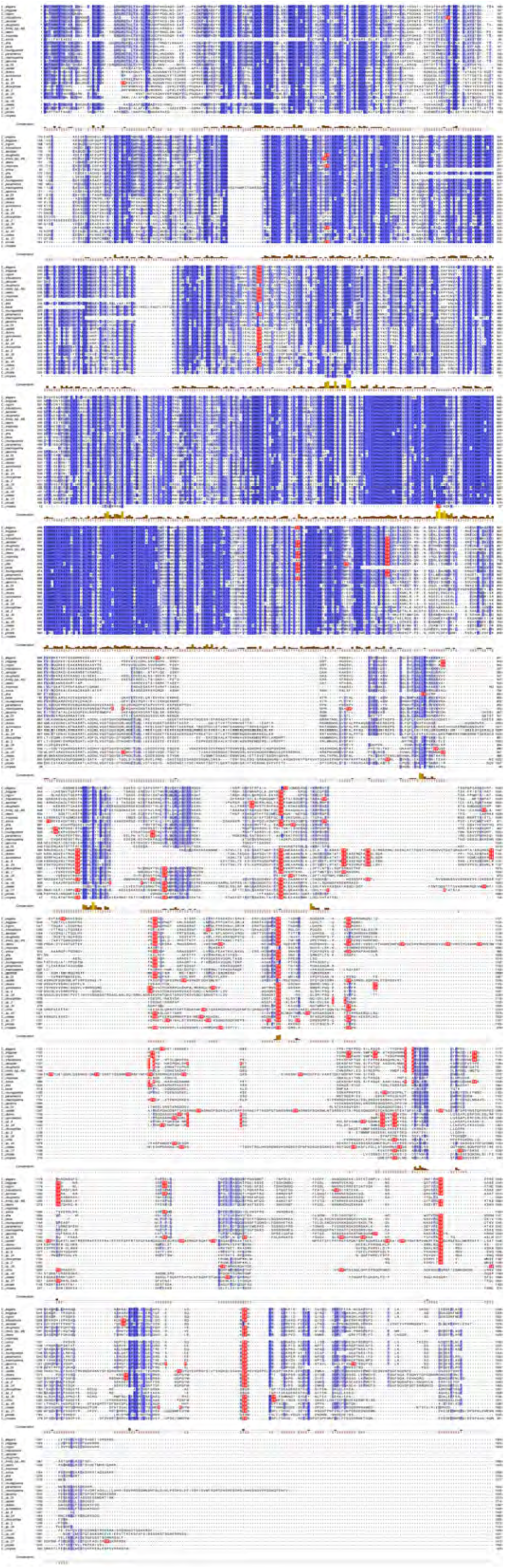
Multiple sequence alignment of MSH-5 sequences from several Caenorhabditid species. TP motifs are highlighted to show their abundance in the C-terminal domains, which are not well conserved and are predicted to be intrinsically disordered.

**Figure S9.**
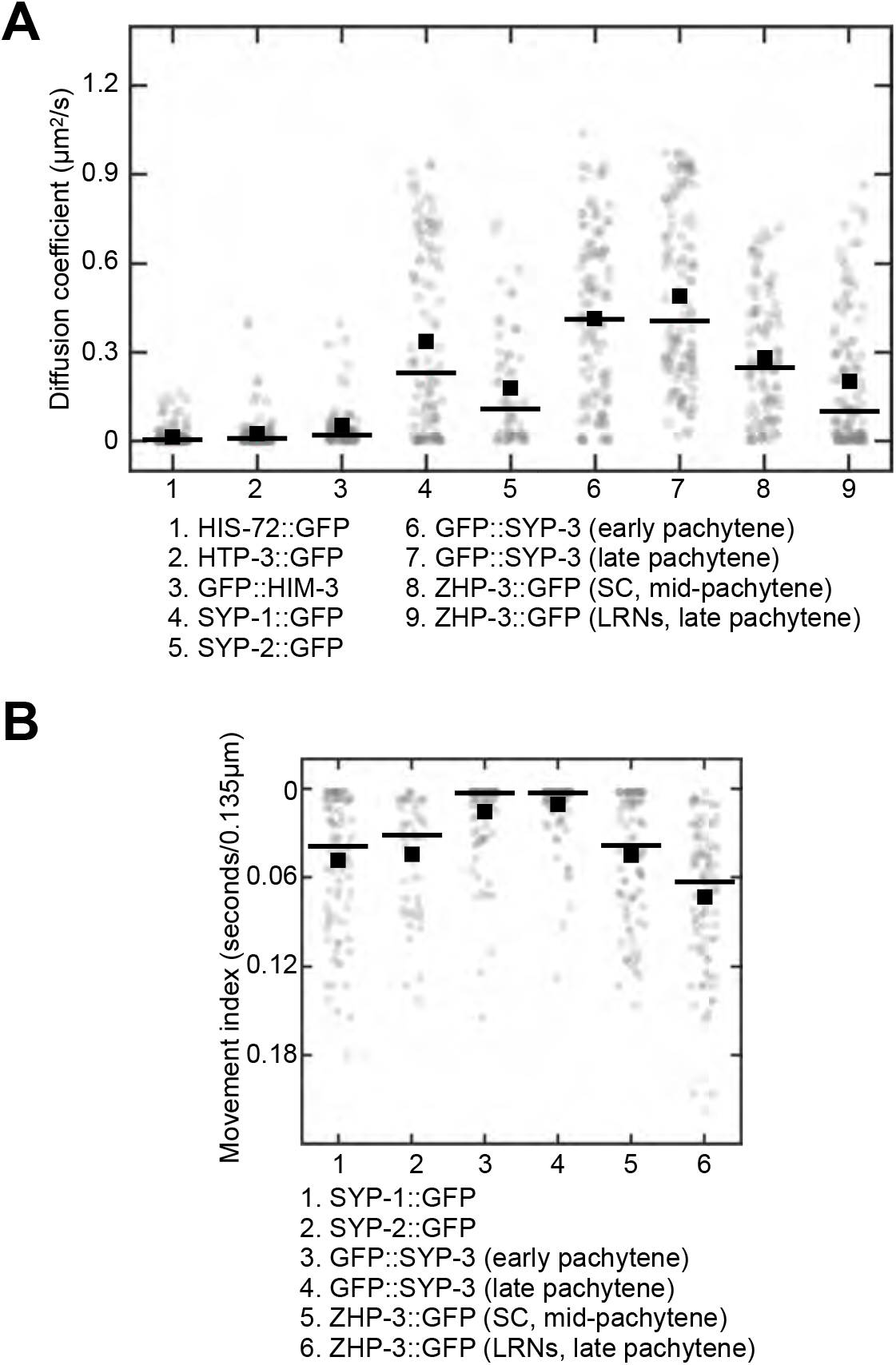
Voxel and voxel-pair based analysis. (A) Diffusion coefficients were derived from analysis of on ACFs based on individual voxels are shown. Black boxes and bars are the average and median diffusion coefficient, respectively. (B) Movement indices for analysis based on pCFs of single voxel pairs are shown. Black boxes and bars are the average and median diffusion coefficient, respectively.

